# The structural mechanism of eukaryotic fluoride channel activation and inhibition by monovalent cations

**DOI:** 10.64898/2026.04.21.719972

**Authors:** Chia-Yu Kang, Sahar Heidari, Sabrina Kolb, Minjun An, Melanie D. Ohi, Hedieh Torabifard, Randy B. Stockbridge

## Abstract

Fluoride is an environmentally prevalent inhibitor of fungi, plants, and other eukaryotes. The fluoride exporter (FEX) is the major resistance mechanism that prevents intracellular fluoride accumulation in eukaryotes. FEX activity is sodium-dependent, but the mechanism for Na^+^ activation and the impact of other cations on FEX function remain poorly understood. Here, we show that sodium and lithium have different effects on channel activity. We leverage these differences to understand how monovalent cations regulate FEX. Functional assays in a reconstituted system show that lithium acts as a competitive antagonist of channel activation by competing with sodium for the central cation binding site. We further demonstrate that FEX inhibition by lithium has consequences *in vivo,* as lithium markedly reduces fluoride tolerance of yeast. A cryo-EM structure of *Candida albicans* FEX with Li^+^, together with molecular dynamics (MD) simulations, reveals the structural mechanism underlying cation-dependent FEX activity. Whereas sodium binding supports a dynamic structure with a broader pore radius, lithium coordination causes a more rigid conformation that is more compact in the channel’s vestibule. These changes perturb the tilt angle of a pore lining helix, and the alter the rotamer of a key phenylalanine in the pore, which together constrict the permeation pathway. In addition to providing general insight into the mechanism of fluoride channel regulation by monovalent cations, these results identify lithium as a previously unknown environmental antagonist of eukaryotic fluoride export and tie cellular fluoride stress tolerance to the abundance of additional ions in the cellular milieu.

## Introduction

Fluoride is broadly distributed throughout the biosphere, from both natural and human sources(1). Because fluoride inhibits key metabolic processes, most plants and microorganisms have evolved cellular responses to fluoride stress(2–4). Fluoride exporters, including the eukaryotic FEX proteins and their bacterial counterparts, the Flucs, contribute to this response by protecting cells from cytoplasmic fluoride accumulation, enabling survival in fluoride-rich environments(5–7). Both Flucs and FEX are highly selective, thermodynamically passive fluoride channels that do not couple fluoride transport to the translocation of any other ion(8, 9). The Flucs are assembled as antiparallel homodimers(10), and the FEX are monomers with two homologous antiparallel domains(11), but the overall channel fold is otherwise highly conserved(9, 12). The third transmembrane helix of each domain has a ∼5 residue break, and the two α-helix breaks cross over each other near the midpoint of the membrane. Structures of Fluc channels first identified a cation bound at this crossover, coordinated by the backbone carbonyl oxygens(12). Later functional studies identified this cation as Na^+^ and showed that it is inserted when the channel folds and that it is not exchangeable on the day timescale(13, 14). However, the introduction of mutations that enable Na^+^ dissociation revealed that Na^+^ occupancy is essential for fluoride permeation in the Flucs(14), and computational studies establish a role for Na^+^ in facilitating fluoride transport(15). Similar to Fluc, FEX binds Na^+^ at breaks in transmembrane helices 3 and 8, and Na^+^ binding is necessary for channel activity(9). Unlike the Flucs, in yeast and plant FEX homologues, the Na^+^ is dissociable in the wildtype proteins with K_d_ values on the order of 0.5 mM(9), which is within a biologically relevant range. A structure of FEX from *Candida albicans* show that the Na^+^ binding niche is accessible to the external solution(9). These observations raise the possibility that the central cation is a physiological gate of channel function.

K^+^ and Li^+^ also modulate FEX channel activity. Potassium has a high K_d_ of 300 mM, and in simulations, K^+^ is only loosely coordinated within the cation binding motif. When bound, K^+^ activates FEX(9). In contrast, Li^+^ disrupts FEX activity, suggesting that subtle differences in cation radius and coordination may impact the adjacent fluoride permeation pathway. However, the inhibition mechanism and the structural basis by which Li⁺ interferes with FEX function, particularly in the context of Na^+^-dependent activation, remains poorly understood.

In this study, we leverage the different effects of sodium and lithium on FEX activity to understand the structural mechanism for FEX regulation by monovalent cations. By combining fluoride efflux assays with structural analysis of FEX from *Candida albicans*, we find that Li⁺ is a competitive antagonist of channel activation by Na⁺. Li^+^ binds to the helical break motif, triggering a subtle conformational change that narrows the pore and prevents fluoride permeation. Molecular dynamics (MD) simulations further indicate that, relative to Na⁺, Li⁺ is more stably coordinated in the cation-binding niche, strengthens electrostatic interactions with the helical break motif, and shifts FEX toward a more rigid conformation with a narrower and less hydrated external vestibule, providing a dynamic basis for its inhibition of fluoride transport. We additionally show that Li^+^ impacts the fluoride resistance of yeast *in vivo*, consistent with its disruption of FEX activity in the reconstituted system. Together, these findings link environmental lithium to reduced fluoride tolerance in eukaryotes, and provide mechanistic insight into the relationship between cation binding and fluoride transport by eukaryotic fluoride channels.

## Results

### Lithium-sodium competition at the cation-binding motif

We previously showed that Li^+^ suppresses fluoride transport by FEX homologues from *Candida albicans* and *Camellia sinensis* (tea plant) in reconstituted systems(9). However, the mechanism of inhibition has not been established. FEX activation requires binding of Na^+^ or, with weak affinity, K^+^, to the central crossover region, and we hypothesized that Li^+^ competes with binding of the activating cation.

To test this, we assessed fluoride export from liposomes by FEX-CA in mixed Li^+^/Na^+^ conditions (**Figure 1A**). Proteoliposomes were reconstituted at three fixed LiCl concentrations (0.1, 0.6, or 2 mM Li^+^) and with varying Na^+^ between 0.1 and 30 mM. All experiments also include 300 mM K^+^, which is required to maintain electroneutrality in liposome flux experiments. Upon initiation of transport, fluoride is released from liposomes that contain an active FEX channel, ultimately reaching a maximum value that reflects the fraction of active protein in the system (9, 16). Because cation exchange is slow relative to the timecourse of the assay, the proportion of active Na^+^-bound protein and inactive Li^+^-bound protein can be determined from this maximum efflux value (see *Methods*).

**Figure 1.**
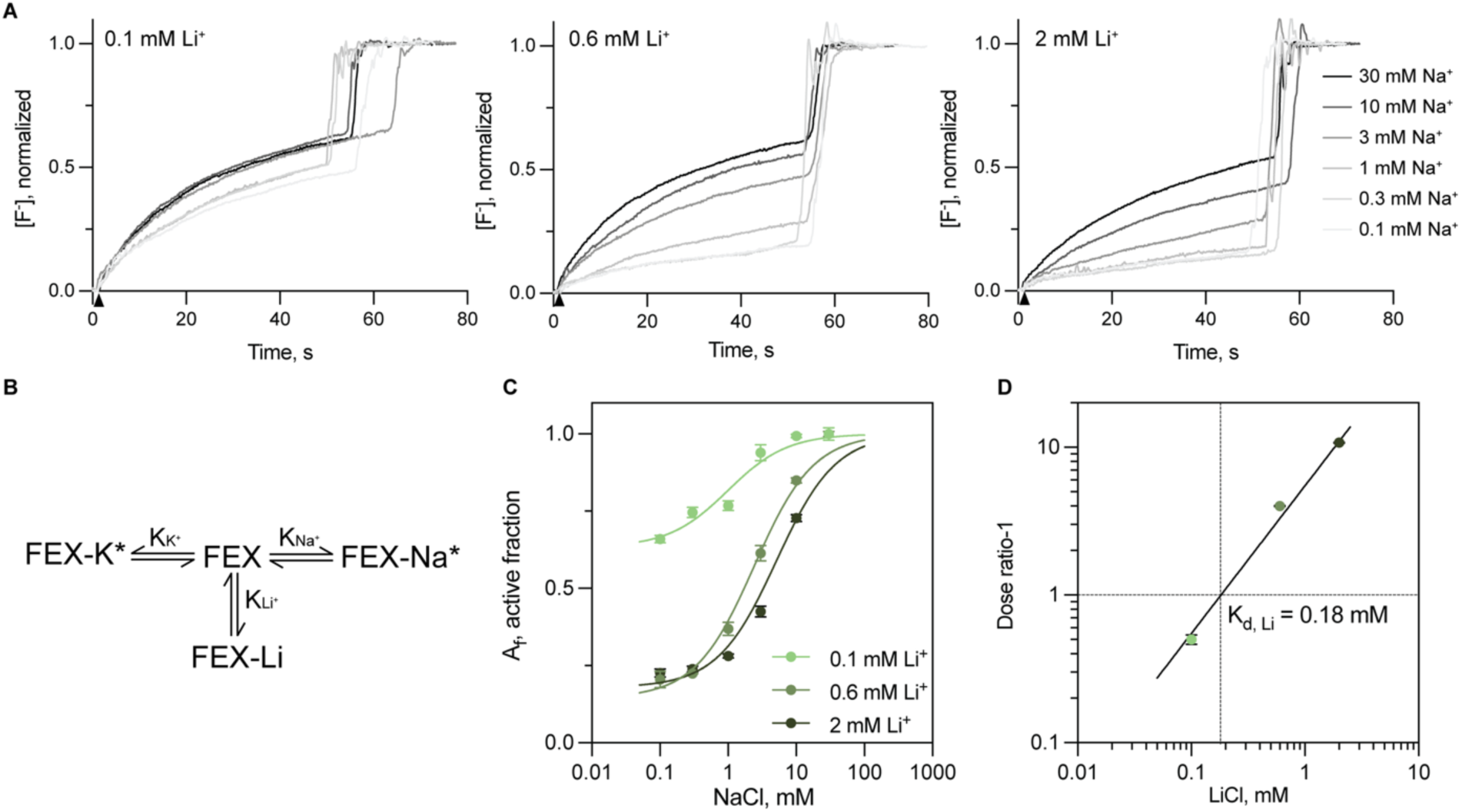
Inhibition of FEX-CA-mediated fluoride efflux by Li^+^. (A) Fluoride efflux measured at three fixed Li^+^ concentrations (left-0.1 mM, middle-0.6 mM, right-2 mM) with increasing Na^+^. Fluoride efflux was monitored using a fluoride-specific electrode. Efflux was initiated by the addition of valinomycin (closed triangle), and traces were normalized against the total encapsulated F^-^ determined after liposome solubilization with N-octyl-β-D-glucoside. Representative traces are shown from three measurements of independent proteoliposome preparations. (B) FEX activation scheme depicting competitive binding of three cations: Na⁺ and K⁺ (agonists), and Li⁺ (antagonist). The asterisk represents the active state of the channel. This scheme assumes that all Na^+^ and K^+^ binding events activate FEX. (C) The fraction of active FEX-CA channels (A_f_) as a function of Na⁺ concentration calculated from fluoride efflux traces, as described in *Methods*. Points and error bars represent the mean and SEM from three independent experiments. Solid lines represent fits to an agonist-response model, yielding K_d_ values for NaCl of 1.0, 2.4, and 5.1 mM at 0.1, 0.6, and 2 mM LiCl, respectively. (D) Schild plot of the apparent K_d_ values determined from the dose-response curves in (C) as a function of the Na^+^ (agonist) to Li^+^ (antagonist) dose ratio. Points are color-coded as in panel C. The solid line shows a fit to a straight line with a slope of 1 (Eq. 5), and the intercept yields a K_d_ of 0.18 mM for Li^+^.

As expected for the scheme shown in **Figure 1B**, the fraction of active protein decreased as Li^+^ concentration increased, with half-maximal channel activation by Na^+^ occurring at 0.6 mM, 2 mM, and 4.7 mM in the presence of 0.1 mM, 0.6 mM, and 2 mM Li^+^, respectively (**Figure 1C**). The Schild plot for lithium is linear with a slope of unity (**Figure 1D**), characteristic of a reversible competitive antagonist (17) and consistent with a model in which lithium, potassium, and sodium compete for the same cation-binding motif. Using previously measured K_d_ values of 300 mM and 0.4 mM for K^+^ and Na^+^, respectively, regression analysis yields a binding affinity of 0.18 mM for Li^+^ (**Figure 1D**). The minimum value for the fraction of active FEX-CA channels, A_f_, determined from each dose response curve is also quantitatively consistent with displacement of K^+^ (held constant in all experiments) by increasing Li^+^. Thus, these experiments provide quantitative evidence that Li^+^ competes with the FEX channel agonist Na^+^, but that Li^+^ binding favors an inactive channel state.

### Lithium reduces the fluoride tolerance of S. cerevisiae in vivo

Since FEX is the primary determinant of fluoride resistance in yeast, its inhibition by Li^+^ suggests that this cation should exacerbate fluoride stress *in vivo*. To test this, we monitored the growth of *S. cerevisiae* in fluoride-containing medium as a function of lithium concentration (**Figure 2**). For wild-type (WT) yeast, the reported minimum inhibitory concentration of fluoride (MIC) is ∼120 mM(5). Consistent with this, 50 mM KF did not impact the growth of WT *S. cerevisiae* in the absence of lithium. However, the addition of LiCl to the fluoride-containing media was associated with significant, dose-dependent reduction in growth, with ∼50% inhibition at 10 mM LiCl and 50 mM KF (**Figure 2A, C**). We confirmed that in the absence of fluoride, lithium causes no detectable growth defect at concentrations up to 100 mM (**Figure 2B, C**), in agreement with prior measurements (18). These results support the idea that Li^+^ is a physiologically important modulator of fluoride tolerance in eukaryotes, in agreement with FEX inhibition by Li^+^ in reconstituted systems.

**Figure 2.**
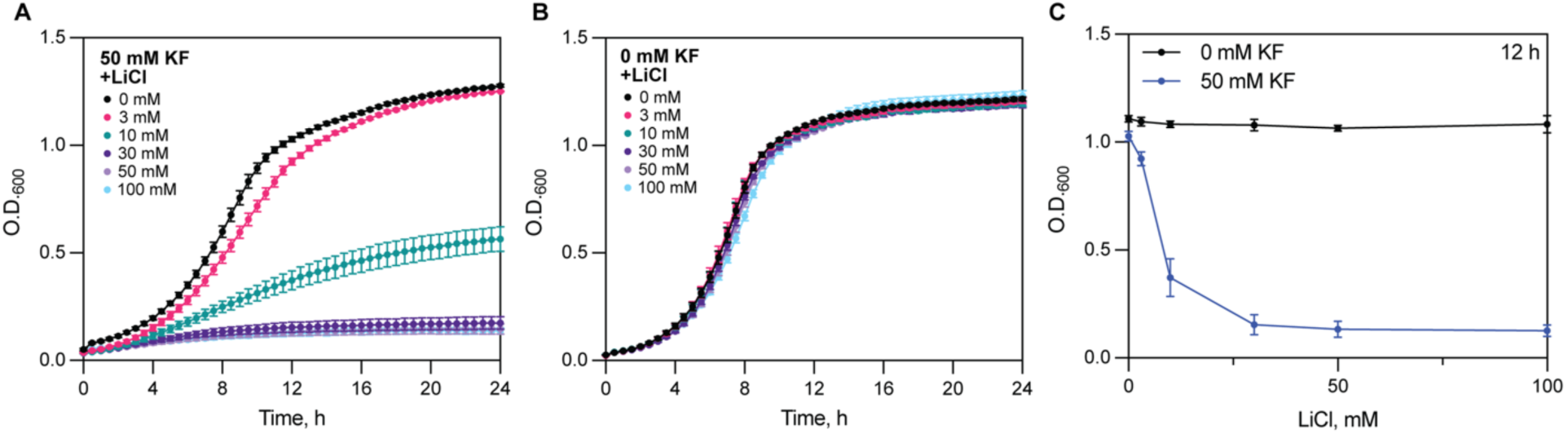
Lithium reduces fluoride resistance in *Saccharomyces cerevisiae*. (A) Growth of wildtype *S. cerevisiae* with 50 mM KF and increasing LiCl as indicated. Cell growth was monitored by measuring optical density at 600nm (OD_600_). (B) Growth of wildtype *S. cerevisiae* grown in in the absence of fluoride and with increasing LiCl. (C) OD_600_ of *S. cerevisiae* at 12 h obtained from growth curves in panels A and B. Points and error bars represent the mean and SEM of four independent biological replicates.

### Cryo-EM structure of Li^+^-FEX-CA exhibits sidechain rearrangement and pore constriction

To understand the structural mechanism of FEX inhibition by Li^+^, we determined a 3.5 Å cryo-EM structure of FEX-CA in the presence of Li^+^ and compared this structure to our previously reported Na^+^-FEX-CA structure(9) (**Figure 3A**). Details of cryo-EM data collection, processing, and structural determination are shown in **Supplementary Figure 1-3 and Supplementary Table 1**). As with the Na^+^-FEX-CA studies, we used a 17-residue N-terminal MPER tag to bind an antibody fragment to serve as a fiducial marker(19). We previously showed that binding of the 10E8v4 Fab fragment to FEX has no impact on channel function(9). To prepare the Li^+^-FEX-CA sample, 150 mM LiCl was added in lieu of NaCl during the final step of protein purification. Prior studies of the cation exchange kinetics indicate that Li^+^ should be the major bound cation under these conditions(9).

**Figure 3.**
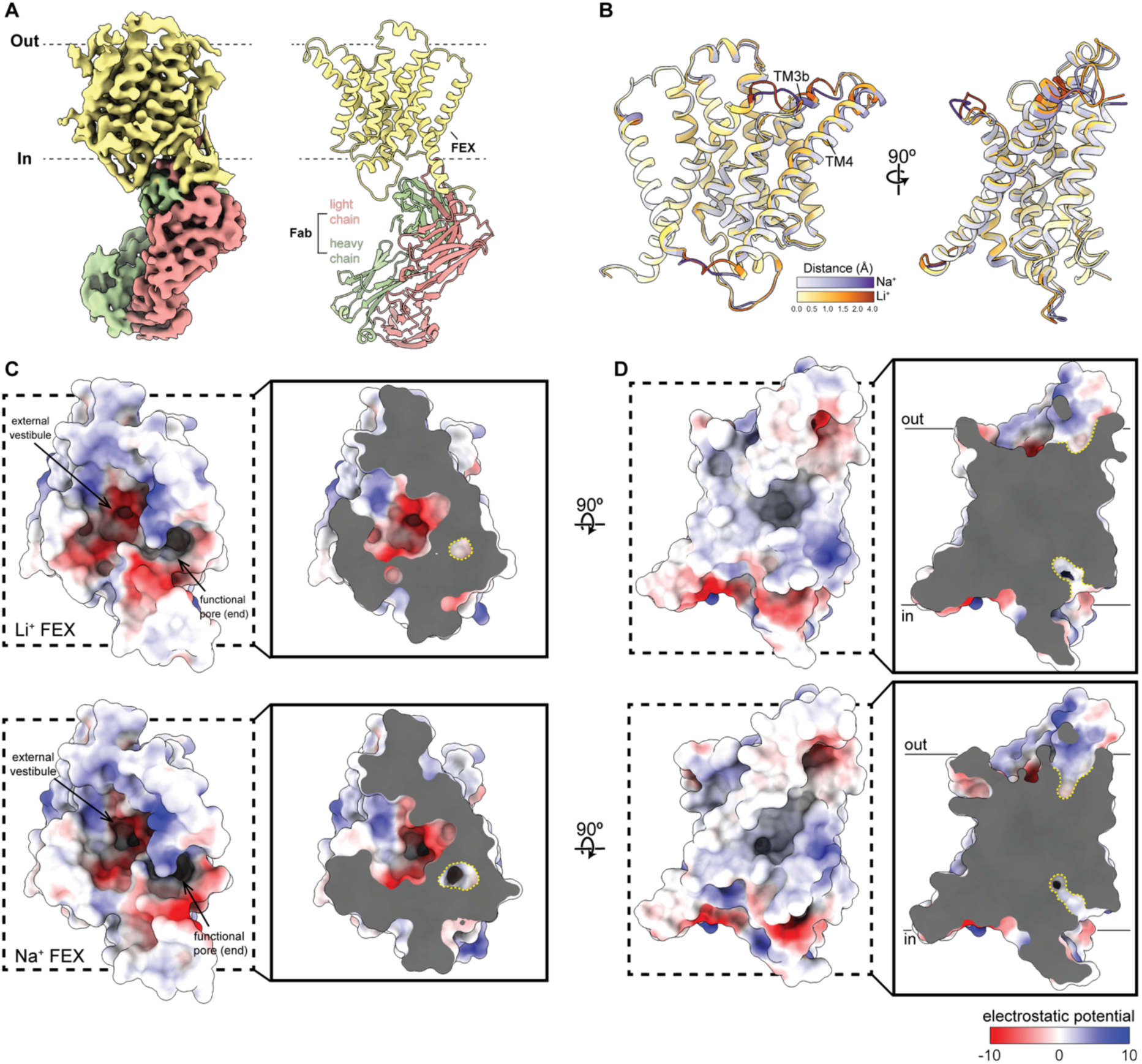
Structure of FEX-CA determined in the presence of Li^+^. (A) Left, cryo-EM map of the FEX-CA/10E8v4 Fab complex. Right, atomic model of FEX-CA/10E8v4 Fab complex. For both panels, FEX-CA is shown in yellow, the 10E8v4 Fab heavy chain in salmon pink, and the light chain in green. Dashed lines indicate the approximate boundaries of the plasma membrane, with the cytoplasmic (in) and extracellular (out) sides labeled. (B) Structural comparison of FEX-CA in the presence of NaCl and LiCl analyzed using *Resi-Ruler*(63). Domain 1 of the Na⁺ and Li⁺ structures are superimposed and the magnitude of the per-residue C_α_ distance differences is displayed as a color gradient (purple, Na^+^-FEX-CA; orange, Li^+^-FEX-CA). (C-D). Cross-sectional views of FEX-CA structures determined with Li⁺ (top) or Na⁺ (bottom). Cavities of the functional pore are shown from the extracellular view (C) or side (D), with cross-sectional regions at the same level, as indicated by yellow dashed lines. Protein surfaces of FEX-CA are colored by electrostatic potential as shown in the scale.

The overall structural differences between Li^+^-FEX-CA and Na^+^-FEX-CA are subtle, with an overall root-mean-square deviation (RMSD) of 1.57 Å over 324 residues. However, despite the two structures sharing many of the same structural features, we observed notable displacements of pore-lining helices TM3b and TM4, along with changes to some loop regions when FEX-CA is bound to Li^+^ (**Figure 3B, Supplementary Video 1**). The pore cavity toward the end of the permeation pathway is shortened in Li⁺-FEX-CA relative to Na⁺-FEX-CA (**Figure 3C-3D**), and the external vestibule becomes more compact in the Li^+^ condition (**Figure 3C**). Since the vestibule shape is defined by the TM3 crossover, this suggests that association with lithium might induce conformational changes within the cation-binding motif.

We next analyzed the fluoride conduction pore of Li^+^-FEX-CA using MOLE(20). Relative to the structure determined with Na^+^, the Li^+^-FEX-CA conduction pathway is more constricted, with the most pronounced narrowing observed along the phenylalanine-lined region near F159, a key residue for channel function(9). The side chain of F159, which is located half a helical turn from the cation-coordinating residue S157, adopts an upward-tilted conformation that does not fit well in Na^+^-FEX-CA map (**Supplementary Figure 4A, 4B**). Real space difference maps calculated between the resolution-matched maps reveal a distinct shift in the density map around F159 (**Supplementary Figure 4C**). This F159 configuration reduces the pore radius to ∼0.4 Å (**Figure 4A-4B**), and might also be less conducive for the edge-anion interactions that characterize fluoride binding and permeation in the Fluc/FEX family(12, 21). A second narrowing of the pore occurs between I163 and F373, forming a second hydrophobic constriction along the pore axis (**Figure 4B**).

**Figure 4.**
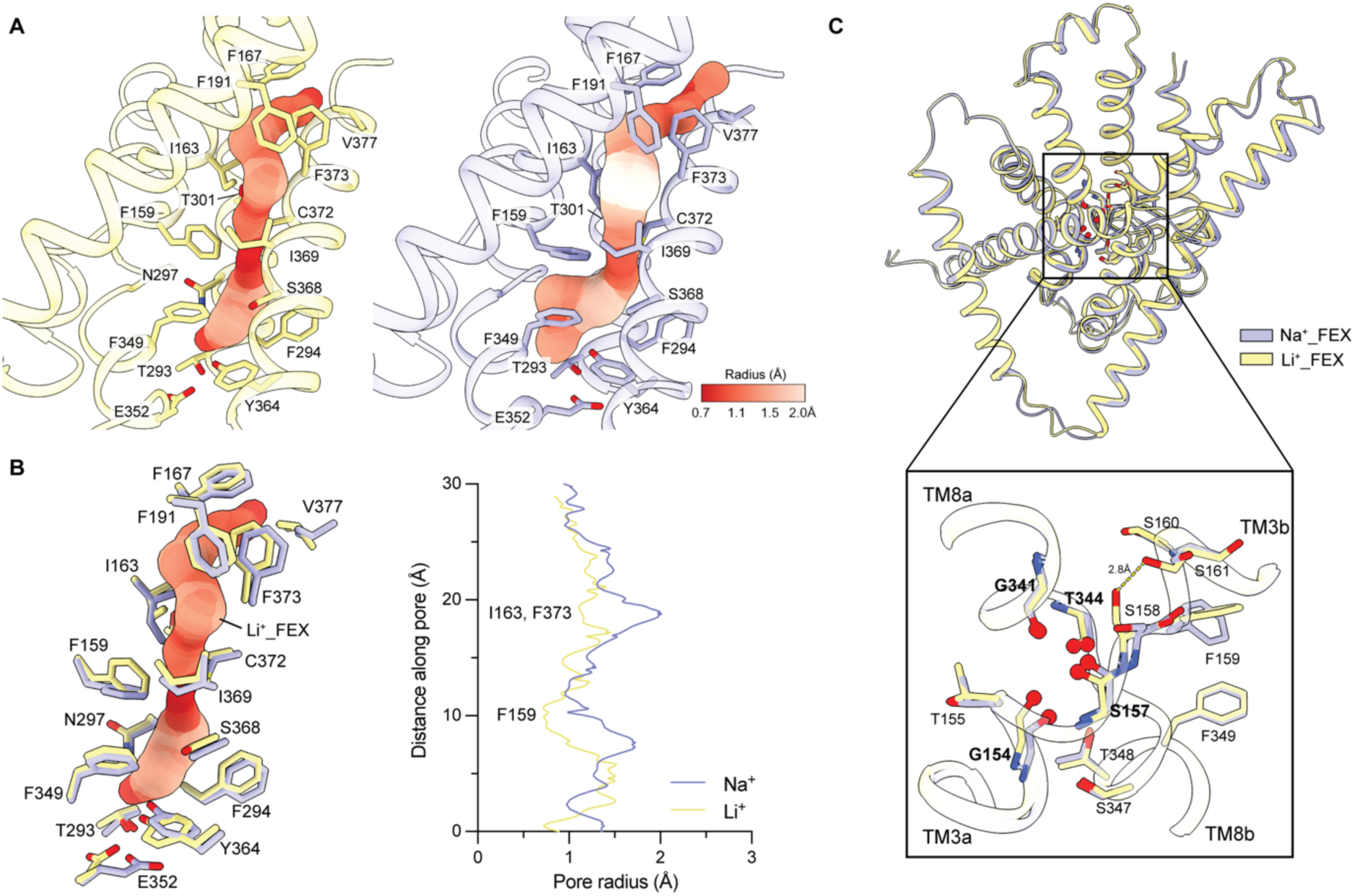
Structural analysis of the fluoride conduction pathway in FEX-CA. (A) Side view of the fluoride conduction pathway identified by MOLE(20) for Li^+^-FEX-CA (yellow) or Na^+^-FEX-CA (purple). For clarity, regions of the protein other than pore-lining TM3b, TM4, TM4, TM8b, and TM9 are omitted. The pore is colored according to its radius, as indicated by the scale. Residues lining the conduction pathway are shown as sticks. (B) Left, the conduction pathway of Li⁺-FEX-CA with surrounding residues shown as yellow sticks. For comparison, the same residues from the Na⁺-FEX-CA structure are overlaid. Right, plot of the pore radius as a function of distance along the pore, corresponding to the view in the left panel, calculated using MOLE. (C) View from the extracellular side of the putative cation-binding motif for Li^+^-FEX-CA (yellow) or Na^+^-FEX-CA (purple). Residues from the TM3-TM8 helical breaks are represented as sticks, with sidechains colored yellow (Li^+^) or purple (Na^+^). Four coordinating carbonyl oxygen atoms proposed to interact with cations are shown as red spheres. In the zoomed-in view, the yellow dashed lines show potential interactions between the side chain of S158 and S161.

The ionic radius of Li^+^ is 0.26 Å smaller than Na^+^, and Li^+^ favors fewer ligands in a tighter coordination sphere. With an atomic mass of just 7, we do not expect to observe lithium density within the cation-binding motif. However, we do find that the four coordinating carbonyl oxygen atoms (**Figure 4C, inset**) proposed to interact with cations adopt a more compact conformation in the Li^+^-FEX-CA structure. The side chain of S158, immediately adjacent to cation-binding carbonyl of S157, also adopts a distinct rotamer, moving away from its interaction partner, R89, in the Na⁺-FEX-CA structure, and to within hydrogen bonding distance of the S161 sidechain in the Li^+^-FEX-CA structure (**Supplementary Figure 4A**, **Supplementary Figure 5**). The vestibule-facing interaction between S158 and S161 perturbs the local backbone geometry of the TM3 break. These residues, which are not constrained by α-helical secondary structure, include pore-lining F159, the sidechain located at the tightest pore constriction. Thus, this structural comparison suggests a mechanism by which subtly altered geometry in the cation binding motif is transmitted via the S158 sidechain to pore-lining residue F159, whose displacement constricts the pore. The FEX-CA mutant F159I was previously shown to impair fluoride resistance in yeast(9). Likewise, S158 is conserved, and, although it faces away from the fluoride conduction pathway, disruption of its packing by mutation to valine and has been shown disrupt FEX activity in the *S. cerevisiae* homolog(22). Thus, prior experimental results support the idea that perturbation of these sidechains can directly modulate channel function.

### MD simulations implicate protein dynamics and sidechain rearrangements in pore constriction

To further substantiate the molecular events associated with cation binding, we performed MD simulations to compare the stability of Li⁺ and Na⁺ in the cation-binding site and to assess the structural consequences of Li⁺ binding for pore architecture. Simulations were initiated from the Na^+^-FEX-CA structure(9) with Li^+^ placed in the cation niche. RMSD analysis (**Figure 5A**) shows that FEX remains more rigid throughout the simulation in the presence of Li⁺ than Na⁺. Although simulations were initiated from the Na^+^-FEX-CA structure, the resulting RMSD distribution of the simulation in the presence of Li^+^ is very similar to the Li^+^-FEX-CA cryo-EM structure, differing only by a small shift (yellow distributions in **Figure 5A**). These results indicate that Li^+^ substitution induces a conformational transition away from the initial Na^+^-bound state toward Li^+^-FEX-CA conformation, while also producing a more rigid conformational ensemble.

**Figure 5.**
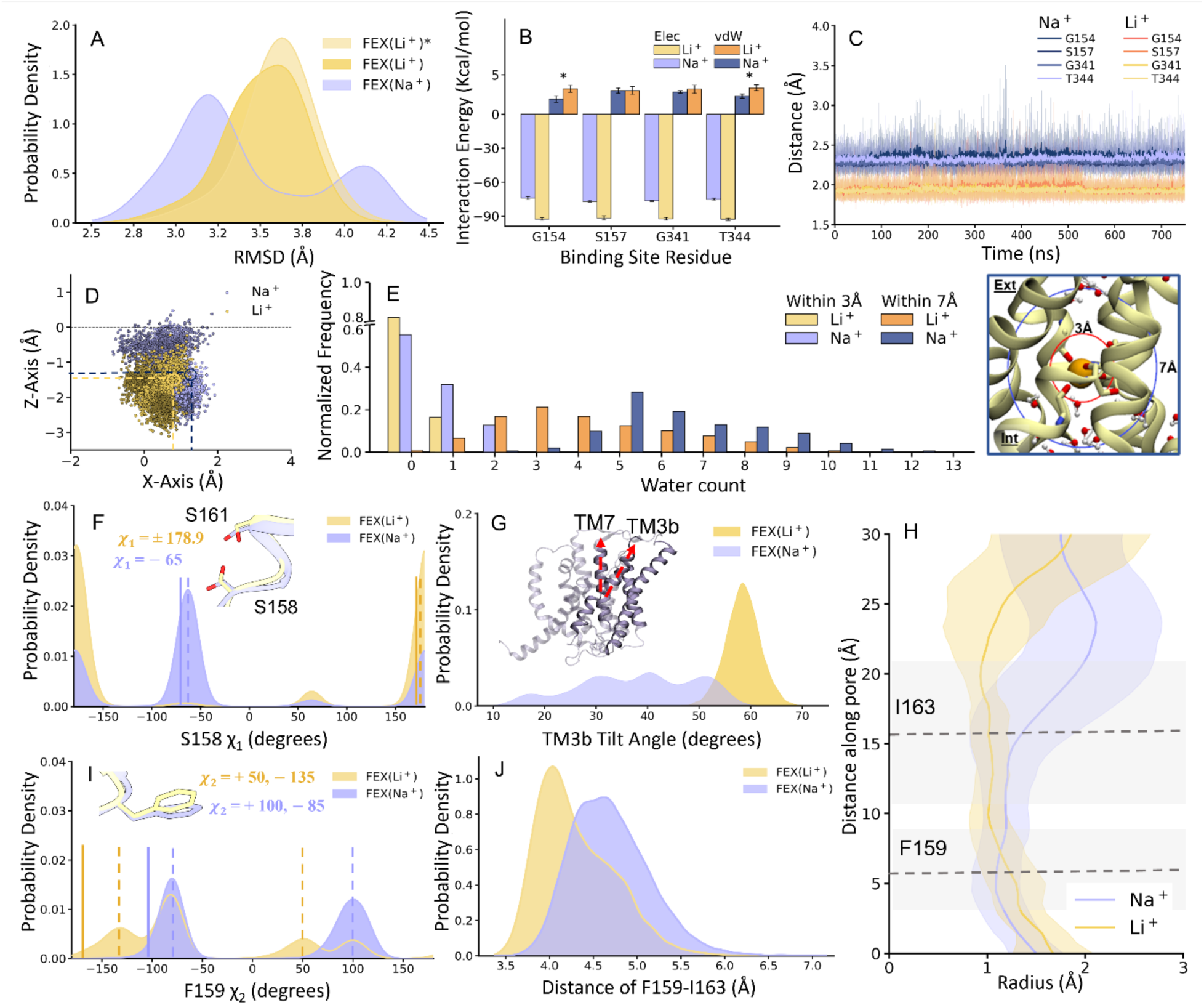
Molecular dynamics analysis of structural differences between Li⁺- and Na⁺-bound FEX-CA. (A) Root mean square deviation (RMSD) of FEX-CA with Na^+^ or Li^+^ placed in the niche over the simulation trajectories relative to the Na^+^-FEX-CA cryo-EM structures. Asterisk (*) indicates the Li^+^-FEX-CA cryo-EM structure used as a reference. (B) Electrostatic (Elec) and van der Waals (vdW) interaction energies between the bound Na⁺ or Li⁺ ion and the cation-binding site residues. Asterisks (*) denote statistically significant differences in vdW interaction energies between the Li^+^- and N^+^-bound states (*P* < 0.05, two-tailed Welch’s *t* test). (C) Distance between the coordinating carbonyl oxygens of the cation-binding site and the Na⁺ or Li⁺ ion as a function of simulation time. Data are represented as the mean (solid line) ± s.d. (shaded area) of three independent replicates for each system. (D) Time evolution of Na⁺ and Li⁺ positions in FEX-CA across all trials, indicated by a color gradient from light to dark. The dashed line marks the center of the bilayer, and negative and positive Z values correspond to the intracellular and extracellular sides, respectively. Small circles mark the approximate initial positions of Na⁺ and Li⁺. (E) Normalized distributions of the number of water molecules within 3 Å and 7 Å of the Na⁺ and Li⁺ ions in the cation-binding niche. (F) χ1 dihedral angle distributions of S158 in Li⁺-FEX-CA and Na⁺-FEX-CA. Dashed lines indicate the χ1 angles corresponding to the most probable S158 orientations sampled in the simulations, which are shown in the inset in stick representation. The solid lines indicate the χ1 angle of S158 in the cryo-EM structures. (G) Tilt-angle of TM3b with respect to TM7 in Li^+^-FEX-CA and Na^+^-FEX-CA structural ensembles. (H) Permeation pathways in Li⁺-FEX-CA and Na⁺-FEX-CA defined using CAVER 3.0 analysis over 2250 trajectory frames for each system. Solid colored lines represent representative pore profiles, and the dashed lines indicate the positions of I163 and F159 along these representative pore profiles. The shaded regions indicate the range spanned by all pores identified in each structural ensemble, and the gray shaded regions denote the range of the I163 and F159 positions across all CAVER-defined pores. (I) χ2 dihedral-angle distributions of F159 in Li⁺-FEX-CA and Na⁺-FEX-CA. Dashed lines indicate the χ2 angles corresponding to the most probable F159 orientations sampled in the simulations, which are shown in stick representation. The solid lines indicate the χ2 angle of F159 in the cryo-EM structures. (J) Distance distribution between the side chains of F159 and I163 in Li⁺-FEX-CA and Na⁺-FEX-CA.

MD simulations indicate that Li⁺ is more stable than Na⁺ in the cation-binding site (**Figures 5B, 5C**), in agreement with K_d_ values measured in liposome efflux experiments. From the simulations, the distance between the bound cation and the four coordinating carbonyl oxygen atoms of the binding motifs was approximately 0.5 Å shorter in Li⁺-FEX-CA than in Na⁺-FEX-CA, indicating stronger coordination of Li⁺ in the niche (**Figure 5C**). Electrostatic and van der Waals (vdW) interaction energies were consistent with this result, showing more favorable electrostatic interactions for Li⁺ together with less favorable vdW interactions, likely reflecting tighter packing and increased short-range repulsion in the Li⁺-bound state (**Figure 5B**). Overall, the larger electrostatic contribution dominates, yielding a more favorable net interaction energy for Li⁺. Tracking cation positions throughout the trajectories likewise showed that Li⁺ remained more tightly confined within the binding site, whereas Na⁺ exhibited greater positional fluctuations relative to the coordinating motifs (**Figure 5D**). Consistent with these differences, Li⁺ remained dehydrated within 3 Å for more than 80% of the simulation time, whereas Na⁺ was water-free only about 55% of the time. In contrast, two water molecules were observed within 3 Å of Na⁺ for roughly 12% of the trajectory, whereas Li⁺ was never coordinated by more than one water molecule, and even that occurred for less than 20% of the simulation time (**Figure 5E**). Together, these results indicate that Li⁺ is intrinsically stable in the niche with a fourfold coordination environment, whereas Na⁺, which prefers a coordination number of six, is more dependent on nearby waters to stabilize its binding.

In line with the increased rigidity of Li⁺-FEX-CA relative to Na⁺-FEX-CA, there are also fewer water molecules within 7 Å of Li⁺ than around Na⁺, reflecting a more compact external vestibule in the Li⁺-bound conformation (**Figure 5E**). Distance and dihedral-angle analyses across the trajectories also support the structural observation of a change in the main rotamer of the vestibule-facing sidechain S158. In Li⁺-FEX-CA, S158 remains closer to S161, while positioned somewhat farther from R89 (**Supplementary Figure 6A, B**). Consistent with these distance distributions, dihedral angle analysis indicates that the S158 rotamer is preferentially oriented toward TM3b residue S161 in the Li⁺-FEX-CA simulation, matching the conformation of the sidechain in the Li^+^-FEX-CA cryo-EM structure. In contrast, in the Na⁺-FEX-CA simulation, the S158 sidechain position fluctuates. S158 interacts with S161 for part of the simulation time, but it more frequently adopts the opposite rotamer, oriented towards the Na^+^, matching the conformation observed in the Na^+^-FEX-CA cryo-EM structure (**Figure 5F**, **Supplementary Figure 7**). In this position, the sidechain oxygen of S158 helps stabilize the Na^+^, including through direct coordination, fulfilling the Na^+^’s preference for more ligands in its coordination sphere.

We next wondered how the greater structural rigidity and local structural changes associated with the Li^+^ occupancy in the niche influenced the pore. Consistent with the cryo-EM structures, analysis of the pore-lining helices TM3b, TM4, TM7, and TM9 further reveals that the main deviations in RMSD between Li⁺-FEX-CA and Na⁺-FEX-CA arise from TM3b and TM4, whereas TM7 and TM9 display similar displacements in both systems (**Supplementary Figure 7**). Root mean square fluctuation (RMSF) analysis further supports this observation by showing reduced fluctuations across Li⁺-FEX-CA residues, except at the end of TM3b and beginning of TM4, particularly in the intervening loop region (**Supplementary Figure 8**), where the conformational change is observed. Tilt-angle analysis across all simulation replicas indicates that TM3b plays a key role in shortening the end of the permeation pathway (**Figure 5G**). In Na⁺-FEX-CA, TM3b exhibits a broader range of motion, with angles spanning ∼10° to 60°. By contrast, in Li⁺-FEX-CA, TM3b is restricted to higher tilt angles, spanning ∼50° to 70°, consistent with a shift toward TM4 that narrows the pore near the end of the permeation pathway in cryo-EM structures (**Figures 5G**).

We further analyzed pore-lining sequences across MD trajectories for each of the Li⁺-FEX-CA and Na⁺-FEX-CA systems using CAVER 3.0(23) (**Figure 5H**). The pore in Li⁺-FEX-CA remains narrower from F159 towards the external vestibule. The greater tilt of TM3b toward TM4 brings the ends of these helices closer together, leading to pore narrowing at the I163/F373 constriction. Fluctuations in radius near F159 likely reflect the changes in the orientation of the F159 side chain during the simulation. Analysis of the F159 χ2 dihedral angle, defined by C_α_, C_β_, C_γ_, and C_δ2_ (**Figure 5I**), consistently indicates that F159 undergoes an approximately 50° rotation in Li⁺-FEX-CA relative to Na⁺-FEX-CA, giving rise to conformations corresponding to the upward-tilted, constricted-pore state. This closely matches the cryo-EM observations, in which the F159 aromatic ring of Li⁺-FEX-CA is rotated by approximately 65° relative to its orientation in Na⁺-FEX-CA, producing a nearly perpendicular arrangement associated with pore narrowing (**Supplementary Figure 4A**). The upward-tilted conformation of F159 allows closer interaction with I163. Consistently, distance analysis of MD trajectories shows that the F159–I163 separation is shorter in Li⁺-FEX-CA than in Na⁺-FEX-CA (**Figure 5J**), while the dihedral angles of I163 and F373 remain similar in both Li^+^- and Na^+^-bound systems (**Supplementary Figures 6C, D**). Thus, the altered F159 orientation in Li⁺-FEX-CA appears to promote TM3b tilting and thereby reinforce narrowing the end of the pore. Together, these structural and dynamic observations support a model in which Li⁺ stabilizes FEX in a constrained, transport-incompetent conformation with a constricted pore. This draws a contrast with the Na^+^-activated FEX structure, which exhibits greater flexibility and samples conformations with a broader pore radius.

## Discussion

In this study, we identify lithium as a new antagonist of FEX channels that diminishes fluoride resistance in cells. Using fluoride efflux measurements, cryo-EM, and molecular dynamics simulations, we establish a mechanism for lithium-dependent inhibition of the eukaryotic fluoride channel from *C. albicans*, FEX-CA. Lithium competes with the agonist sodium at a shared cation-binding motif, exhibiting 2-3 fold higher affinity, but its binding induces a transport-inactive conformation.

Prior studies have shown that Na^+^ occupancy is important for function in both FEX(9) and the structurally related bacterial Flucs(13, 14), but the mechanism of channel activation by cations has not been established. Cation-free structures of fluoride channels have not yet been attained, but the structural rearrangements coincident with Li^+^ binding provide insight into cation-dependent pore opening (**Figure 6**). Both our structures and simulations converge on the finding that Li^+^ adopts a tighter, and more compact coordination geometry than Na^+^ within the cation niche motif. MD simulations suggest that Li^+^ bound FEX is overall less dynamic than the Na^+^ bound protein, which might explain the better structural resolution achieved in the presence of Li^+^ compared to Na^+^. Simulations also suggest that when the Li^+^ occupies the niche, its preference for fewer coordinating ligands means that vestibule sidechain S158 is not necessary for cation stabilization, and also that fewer waters approach the cation over the course of the simulation. These small changes in the vestibule’s hydration and the cation’s immediate coordination sphere are accompanied by rotamerization of S158, prompting an interaction with TM3b sidechain S161 that causes local rearrangements and fluctuations in TM3b that constrict the pore radius. Both the cryo-EM maps and the MD simulations implicate F159 in this response, as its side-chain orientation is consistent with the TM3b shift associated with pore narrowing. In contrast to most anion channels, which tend to exhibit broader anion selectivity, fluoride channels achieve high selectivity over biological competitor Cl^-^(7, 8), which likely contributes to the channel’s sensitivity to the pore restriction.

**Figure 6.**
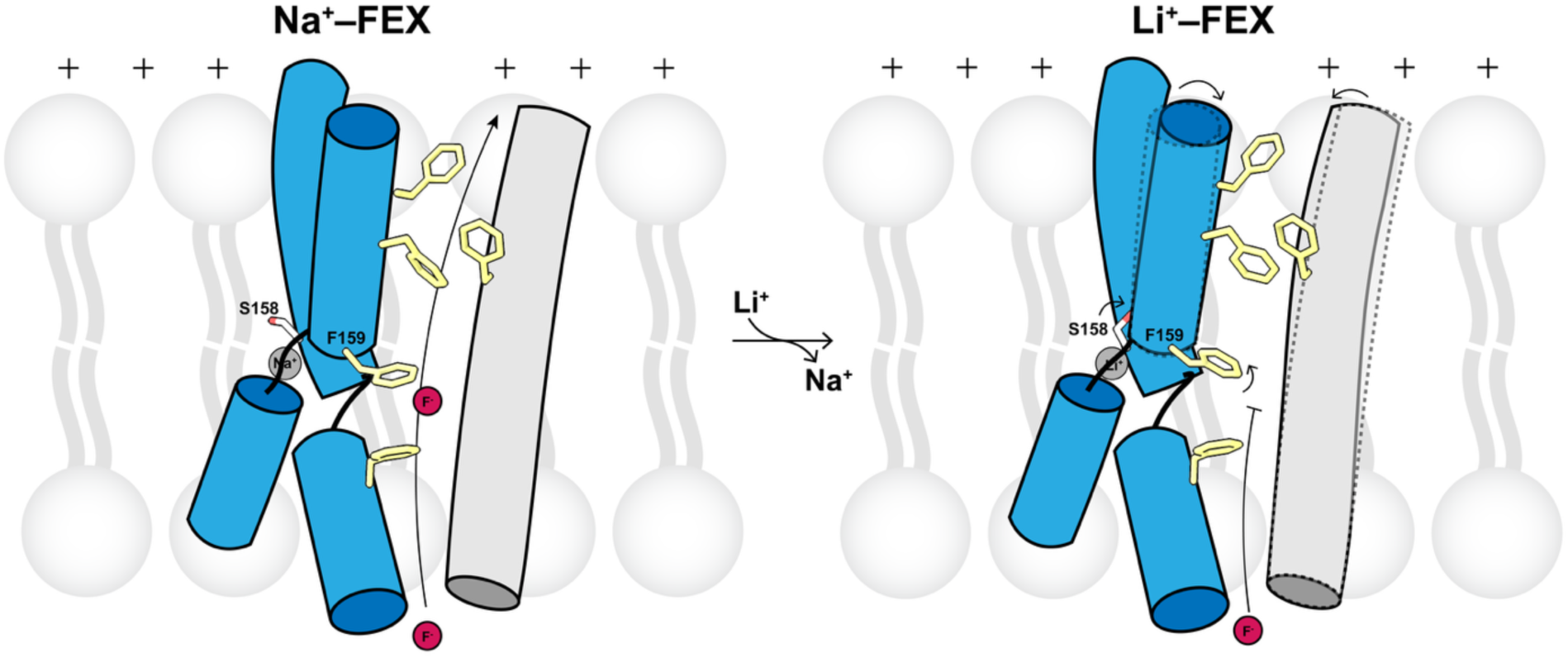
Model of cation-dependent channel opening in FEX-CA. In this model, changes to the coordination sphere of the central cation are broadcast to the pore via a rotameric change in S158. This causes displacements of helices TM3b and 4 (dashed lines), and a change in the favored rotamer of F159, which together constrict the pore.

Although Li^+^ has not been previously recognized as a FEX channel inhibitor, this cation is known to disrupt the activity of a range of Na^+^-coupled transporters(24–27). Due to its small ionic radius, high polarizing strength, and low coordination number, replacement of biologically relevant cations by Li^+^ often fails to preserve biological function in Na^+^-dependent enzymes and transporters (28–30). In FEX, K^+^ can also access this conserved binding site, but it is also a poor replacement for Na^+^. K^+^ is only loosely bound to the site, with a K_d_ value is >750-fold greater than that of Na^+^, and simulations show that K^+^ is more extensively hydrated and has a higher propensity to leave cation binding site(9). The variable effects of Li^+^, Na^+^, and K^+^ on FEX function demonstrates the importance of cation coordination geometry in supporting the open structure of FEX, and establishes that activation of FEX by Na^+^ is not due solely to electrostatic stabilization of the fluorides, but that the Na^+^ plays a mechanistic role, as in the Flucs(15).

Finally, the yeast resistance assays described here establish Li^+^ as a relevant *in vivo* modulator of cellular fluoride resistance. Although Na^+^ is more biologically abundant, many natural environments contain appreciable Li^+^ as well(31). Li^+^ is also a prominent anthropogenic environmental pollutant(31). Various mechanisms of Li^+^ toxicity in yeast have been described, including disruption of ion homeostasis replacement of metal ion cofactors (32–35), and inhibition of central carbon metabolism (34–36). Similarly, in plants, high lithium accumulation causes growth defects, perturbs nutrient uptake, and interferes with cation-dependent signaling pathways(29, 37). Our work establishes a new mechanism of lithium toxicity, the disruption of fluoride stress tolerance. Given the functional similarities between yeast and plant FEX(7), including the dependence on cation binding to the central binding site(9), it is likely that Li^+^ inhibition is also a relevant factor in plant fluoride tolerance, and for other eukaryotes with FEX channels. For such organisms, our results provide a mechanistic foundation to understand the relationship between environmental chemistry and fluoride stress tolerance.

## Materials and Methods

### Yeast strains and growth assay

*S. cerevisiae* wild-type strain BY4741 (*MATa his3Δ1 leu2Δ0 met15Δ0 ura3Δ0*) (Horizon^TM^ Discovery, Lafayette, CO) (38) was grown overnight in 5 mL of YPD medium. The next day, the cells were diluted to an OD_600_ of ∼0.2 in fresh medium and allowed to grow for 5 hours to reach an OD_600_ of 1. The cells were further inoculated into YPD medium at an OD_600_ of ∼0.1 in a 48-well plate, containing LiCl and KF as indicated, and incubated with continuous shaking at 30 °C. Optical density measurements were taken every 30 min for 24 h using a Spark Cyto multimode microplate reader (Tecan, Woburn, MA). YPD was prepared from a commercial yeast extract with minimal residual Na^+^ according to the manufacturer’s analysis (Sigma-Aldrich).

### FEX expression, purification, and reconstitution

Protein expression and purification were carried out exactly as described previously(9). In brief, the gene encoding *Candida albicans* FEX (FEX-CA) was constructed with the pre-signaling peptide of ⍺-mating factor fused to the N-terminus to improve the maturated protein insertion into the plasma membrane, and the 17 amino acid MPER epitope to bind to the 10E8v4 antibody fragment (9, 39). Protein was expressed in *S. cerevisiae* strain BJ2168. Yeast cell pellets were lysed using Zymolase 20T digestion (Amsbio, Cambridge, MA) prior to Dounce homogenization in breaking buffer (20 mM Tris-HCl, pH 8.0, 100 mM NaCl, 5 mM NaF). Lysate was extracted with 2% n-Dodecyl-β-D-Maltoside (DDM) (Anatrace, Maumee, OH) for 2 hours at room temperature and protein purified using Strep-Tactin XT 4Flow resin (IBA Lifescience, Pittsburgh, PA) equilibrated with wash buffer (20 mM Tris-HCl, pH 8.0, 100 mM NaCl, 5 mM NaF, 1 mM DDM) followed by size-exclusion chromatography (SEC) (Superdex 200, Cytiva, Marlborough, MA) in SEC buffer (20 mM 4-(2-Hydroxyethyl)piperazine-1-ethanesulfonic acid (HEPES) pH 7.5, 150 mM LiCl, 5 mM NaF, and 0.5 mM DDM for cryo-EM or 1 mM DDM for functional studies.

Freshly purified FEX-CA was reconstituted with *E. coli* polar lipids (EPL; Avanti Polar Lipids, Alabaster, AL) at a protein:lipid ratio of 1 µg protein/mg EPL (protein:lipid molar ratio of ∼1:50000), solubilized with 35 mM 3-[(3-Cholamidopropyl)-Dimethylammonio]-1-Propane Sulfonate (CHAPS; Anatrace, Maumee, OH). Detergent was removed by dialysis against 4-6 L of reconstitution buffer (typically 15 mM HEPES pH 7.5, 300 mM KF, and indicated concentrations of NaCl and LiCl) over 1.5 days. For cation competition experiments, reconstitution buffer was prepared with 0.1, 0.6, or 2 mM LiCl, with additional NaCl added at varying concentrations (0.1, 0.3, 1, 3, 10, and 30 mM) for each LiCl condition. Following reconstitution, protein/lipid mixtures were subjected to three freeze/thaw cycles and extruded through a 400 nm filter (Cytiva, Marlborough, MA) to generate unilamellar proteoliposomes.

### Fluoride efflux assay

Fluoride efflux from FEX proteoliposomes was measured using a lanthanum fluoride-specific electrode (Cole-Parmer, Vernon Hills, IL) as previously described (8, 40). Proteoliposomes were passed over a Sephadex G50 resin column equilibrated in assay buffer (10 mM HEPES-KOH at pH of 7.5, 300 mM K^+^ isethionate, with LiCl or NaCl as indicated), then diluted 40-fold in assay buffer. Valinomycin (Sigma-Aldrich) was added to a final concentration of 1.8 µM to initiate fluoride efflux. At the end of the timecourse, 30 mM n-octyl-β-D-glucoside (β-OG) (Anatrace, Maumee, OH) was added to disrupt the proteoliposomes and release any remaining encapsulated fluoride. Raw traces were normalized to total encapsulated fluoride, and the fraction of occupied liposomes was converted to the fraction of active protein A_f_ as previously described (9, 16):

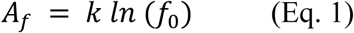

For the competitive agonist/antagonist model shown in **Figure 1C**, A_f_ is equal to:

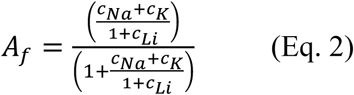

where *c_ion_* is given by:

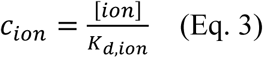

This model assumes that all Na^+^ binding events render the channel active. All experiments were conducted with 300 mM K^+^, equal to FEX-CA’s experimentally measured K_d_ value for K^+^(9), allowing c_K_ to be set equal to one. K_d,Na_ was set to the experimentally measured value K_d_ value for Na^+^ binding to FEX-CA (0.4 mM(9)). For each dose-response curve, A_f_ as a function of Na^+^ was fit to a single-site binding function to extract apparent K_d_ values, K_d, app_, which were expressed as a dose ratio, r:

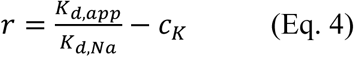

The relationship between the dose ratio and the Li^+^ concentration was determined by Schild regression analysis(17):

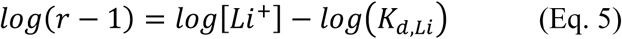

The A_f,min_ values from the fits to the dose-response curve agree with the expected values given by:

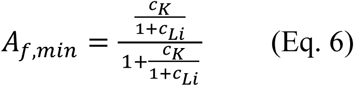

### Fab preparation

10E8v4 antibody expression vectors were obtained from the AIDS Reagent Program, Division of AIDS, NIAID, NIH from Dr. Peter Kwong. Heavy and light chain DNA was transfected into ExpiCHO-S and expressed according to the manufacturer’s protocol (Thermo Fisher Scientific). Six days post-transfection, IgG-10E8v4 was purified as described previously(9, 41). The 10E8v4 Fab fragment was generated by papain digestion (Sigma-Aldrich 1:200, v/v) in phosphate-buffered saline with 1 mM EDTA and 6.5 mM cysteine. The digestion mixture was applied to a protein A affinity column to separate the Fc domain and Fab fragment, followed by SEC (Superdex 200 equilibrated with 20 mM HEPES pH 7.5, 150 mM NaCl, 5 mM NaF).

### Cryo-EM grid preparation and data collection

The MPER-tagged FEX-CA/10E8v4 Fab complex was incubated at a molar ratio of 1:1 for 1-2 hr at room temperature, followed by further purification using SEC (Superdex 200, 20 mM HEPES pH 7.5, 150 mM LiCl, 5 mM NaF, 0.5 mM DDM). After concentrating the complex to 3-4 mg/mL, 3.5 µL of the sample was applied to glow-discharged copper Quantifoil R1.2/1.3 300 mesh grids (Electron Microscopy Sciences, Hatfield, PA, USA). Grids were blotted for 2.5 s at 18 °C, 100% humidity, and plunge-frozen in liquid nitrogen cooled liquid ethane using a Vitrobot Mark IV (Thermo Fisher Scientific). Micrographs were collected at a magnification of ×105,000 (0.834 Å per pixel) using a Titan Krios G3 electron microscope (Thermo Fisher Scientific) operated at 300 kV. The microscope was equipped with Gatan K3 direct electron detector and a Bioquantum Imaging Filter (slit width 20 eV), with a defocus range from -0.8 to -2.5 µm using SerialEM v4.1.1(42). Each image stack was composed of 50 individual frames with a total dose of 50 e^-^ per Å^2^ (**Table 1**).

### Cryo-EM image processing

5,665 images were pre-processed using patch-motion correction, patch CTF correction, and micrograph denoising in cryoSPARC v4.5.1(43, 44), as shown in **Supplementary Figure 2**. 3,566,437 particles were picked using template picker (**Supplementary Figure 2** -dashed line box) with template references that correspond to the FEX-Fab complex, Fab alone, FEX alone, and a DDM micelle, as described previously(9). Particles were extracted using a box size of 320 pixels (0.834 Å/pixel) and were subjected to heterogeneous refinement with the four 3D template volumes (**Supplementary Figure 2** -blue box) to isolate complex particles. The class representing the FEX-CA/10E8v4 Fab complex (660,648 particles) was subjected to 4-class *ab initio* reconstruction followed by heterogeneous refinement. Particles (409,300) in the 3D volumes showing a FEX-CA/10E8v4 Fab complex were then subjected to additional rounds of iterative 2D classification and 3 rounds of 2-class *ab initio* reconstruction followed by heterogeneous refinement. 263,216 particles representing the intact complex were then re-extracted with a larger box size of 352 pixels (0.834 Å/pixel) for 3D classification designating four classes, followed by heterogeneous refinement to separate heterogeneous states and isolate a high-quality complex class that contained 112, 190 particles. Non-uniform refinement was used resulting in a final 3.56 Å resolution map.

### Model building, refinement, and structural validation

The structure of the FEX-CA /10E8v4 Fab determined in Na^+^ (PDB: 9OP1)(9) was used as an initial model to fit the experimental map, followed by manual adjustment in Coot (v0.9.8.93)(45) and real-space refinement in PHENIX (v1.21)(46). Structural validation was performed using PHENIX and MolProbity(47). PyMOL v3.1 (Schrödinger) and ChimeraX v1.8 (48) were used for structural visualization and figure generation.

### Model preparation for MD simulations

Model preparation was carried out as described previously(9). Briefly, the Na⁺-FEX-CA starting structure (PDB: 9OP1) was aligned with Fluc-Bpe (PDB: 5NKQ) to place the Na^+^ ion in the niche. Protonation states of titratable residues at pH 7.4 were assigned using the H++ web server(49, 50). The protein was embedded in a 3:1 POPE/POPG bilayer, solvated with TIP3P(51) water, and neutralized with counterions using Packmol-Memgen(52). NaCl ions were subsequently added to obtain a physiological salt concentration of 0.150 M. The Li⁺-bound structure was modeled starting from the Na⁺-FEX-CA cryo-EM structure to define the transition and structural changes associated with cation replacement. The Li^+^-bound FEX-CA structure was generated by replacing the niche Na^+^ ion with Li^+^ in the prepared FEX-CA structure from the previous work(9) and solvated in LiCl ions. System coordinates and parameters were generated in the tleap module from the AmberTools20(53) using the Amber ff14SB(54), lipid 21(55), and TIP3P(51) force fields. The Li⁺-bound FEX-CA system contained 98,092 atoms in a 105Å × 105Å × 104Å solvated box with approximately 17,758 water molecules.

### Simulation details and analysis

Simulations of the Na⁺-bound FEX-CA system were performed previously and were reported in our earlier study(9). The Na⁺-bound trajectories analyzed here were taken from this previous study and were not regenerated for this work. Three independent production runs were performed for the Li⁺-bound FEX-CA system with pmemd.cuda in Amber20(53, 56). Prior to production, the prepared systems were subjected to sequential minimization, heating, and equilibration steps. The minimization procedure consisted of three stages. In the first stage, only water molecules and hydrogen atoms were allowed to move, while the remaining atoms were restrained with a force constant of 7.5 kcal/mol/Å^2^. In the second stage, ions and protein side chains were also included in the relaxation, with restraints maintained on the protein backbone. In the final stage, all restraints were lifted. Each of the first two minimization stages included 5,000 steps of steepest descent followed by 5,000 steps of conjugate gradient minimization, whereas the last stage used 10,000 steps of each method.

The minimized systems were then heated in two successive intervals, first from 0 to 100 K over 4 ns and then from 100 to 303 K over 8 ns. A Langevin thermostat(57) was used during both heating phases, and the protein and lipids were restrained with a force constant of 5 kcal/mol/Å^2^. Pressure coupling was introduced during the second heating stage using the Berendsen barostat(58). To ensure stability of the periodic boundary conditions, the systems were subsequently equilibrated for ten 0.5 ns intervals before entering production. Each production trajectory was extended to 750 ns under NPT conditions at 303 K, with temperature maintained by the Langevin thermostat(57) and pressure controlled by the Berendsen barostat(58). A 2 fs time step was used, bonds involving hydrogen atoms were constrained with SHAKE(59), and a 12 Å cutoff was applied for nonbonded interactions. Frames were saved every 10 ps. Simulation inputs and parameter set for all systems have been deposited in Zenodo https://doi.org/10.5281/zenodo.19463044.

All production trajectories were analyzed at 100 ps intervals using the cpptraj module in AmberTools20(60) and VMD(61). Root mean square displacement (RMSD) of backbone atoms and root mean square fluctuation (RMSF) of the whole residue were calculated, with respect to the cryo-EM structures of Li^+^-FEX-CA and Na^+^-FEX-CA, as well as the average structures, to monitor overall structural stability and residue-level flexibility. To assess the stability of the Na^+^ and Li^+^ ions in the binding niche, we quantified the number of surrounding water molecules within 3 Å and 7 Å, representing the first and second coordination shells, respectively(62). Electrostatic and van der Waals (vdW) interaction energies between the cation-binding motifs and the Na^+^ and Li^+^ ions were calculated using linear interaction energy (lie) analysis. For each residue, the interaction energies were averaged over independent simulations. The resulting vdW replicate means for the Li^+^- and Na^+^-bound states were compared using a two-tailed Welch’s t-test. Differences with P < 0.05 were considered significant. Distance analysis was also performed to measure the distance between oxygen atoms of cation–binding motifs and the Na^+^ and Li^+^ ions. Moreover, the position of the cations within the binding site were monitored throughout the simulations. To characterize structural changes in the Na^+^- and Li^+^-bound FEX-CA systems, particularly those related to pore geometry, distance, tilt-angle, side chain dihedral-angles, and Caver(23) analyses were performed. Distance analyses were performed between the side-chain centers of mass of F159 and I163, the O_γ_ atoms of S158 and S161, and the O_γ_ atom of S158 and the center of mass of the three guanidinium nitrogen atoms of R89. The TM3b tilt angle was determined by defining two vectors along TM3b and TM7. CAVER 3.0(23) analysis was performed on 2250 frames from all production trajectories of each of the Na^+^- and Li^+^-bound FEX-CA systems. VMD(61) and ChimeraX(48) were also used for visualization of trajectories and generation of structural figures.

## Acknowledgements

This work was supported by National Institutes of Health grants R35 GM128768 (NIH/NIGMS) to R.B.S., S10OD030275 to M.D.O., and R35 GM155106 to H.T (NIH/NIGMS). C.-Y. K. was supported by American Heart Association predoctoral fellowship 24PRE1192512. S.K. was supported by a Horizons Summer Intern Fellowship from the Department of Molecular, Cellular, and Developmental Biology at the University of Michigan. We thank the U-M Cryo-EM facility, which is supported by the U-M Life Sciences Institute, the U-M Biosciences Initiative, and the Beckman Foundation, for scientific and technical assistance, as well as the Office of Information Technology and the Cyber Infrastructure Research Computing (CIRC) at the University of Texas at Dallas and the Texas Advanced Computing Center (TACC) at the University of Texas at Austin for providing HPC resources.

## Author Contributions

C.-Y. K.: conceptualization, investigation, writing – original draft, visualization; S. H.: conceptualization, investigation, data visualization, writing – original draft; M. A.: investigation; S. K.: investigation; M. D. O.: supervision, writing – review and editing; H.T.: conceptualization, data visualization, writing – review and editing, supervision, funding acquisition, project administration; R. B. S.: conceptualization, data visualization, writing – review and editing, supervision, funding acquisition, project administration. The authors declare no competing interests.

## Data and Materials Availability

Coordinates and maps have been deposited in EMDB with PDB ID 11WR and EMD-76140. Simulation input and parameter files for all the systems are available at Zenodo: https://doi.org/10.5281/zenodo.19463044. All other data needed to evaluate the conclusions in the paper are present in the paper or the supplementary materials. FEX expression plasmids are available upon request.

---

**Supplementary Figure 1.**
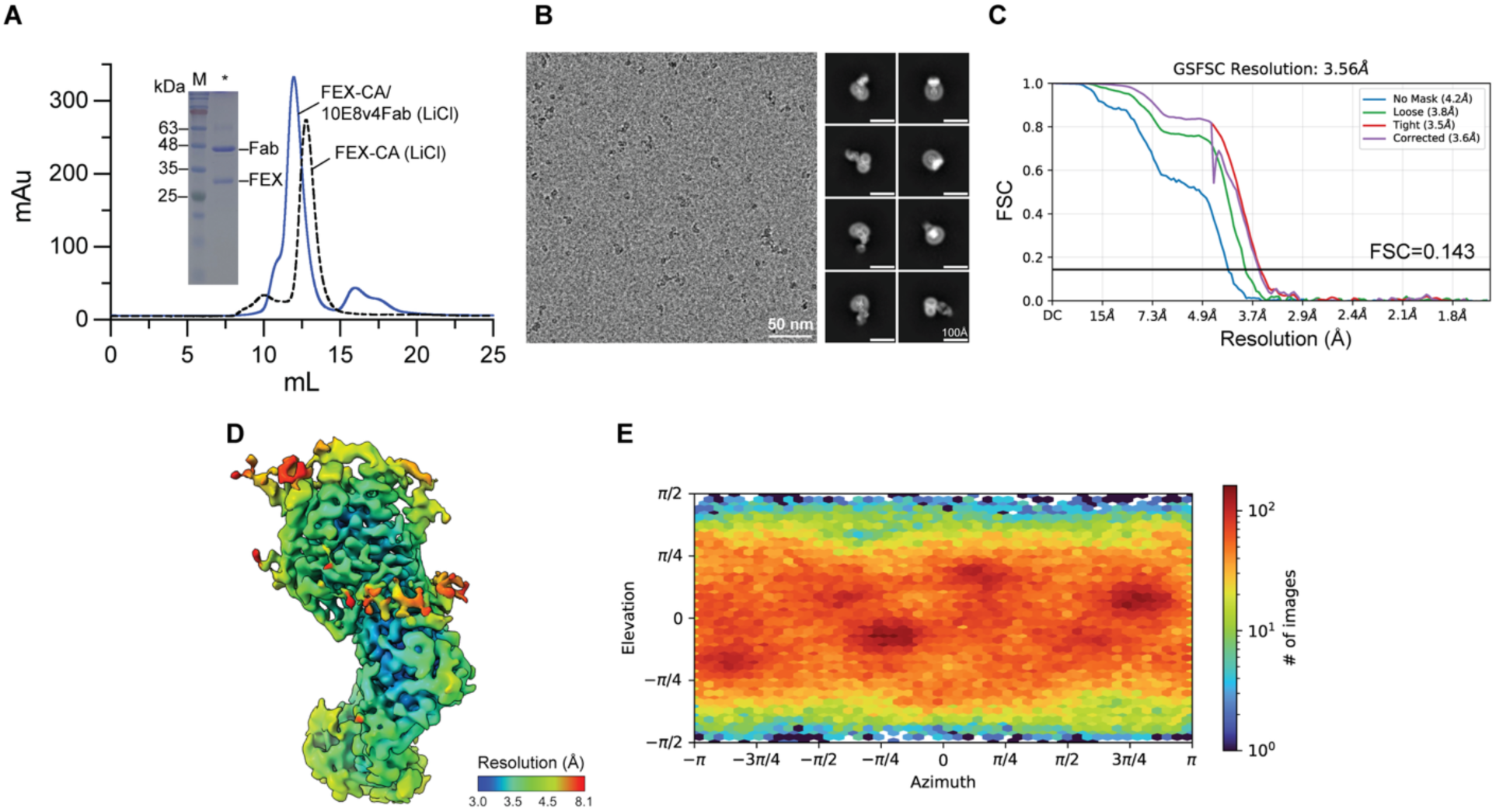
Preparation, micrograph and structure validation of FEX-CA in LiCl. (A) Size exclusion chromatograms of MPER-tagged FEX-CA alone (gray dashed line) or with 10E8v4 Fab at a 1:1 molar ratio (blue trace) in the presence of 150 mM LiCl. Inset: Coomassie-stained SDS-PAGE of the peak fraction corresponding to the FEX-CA/10E8v4 Fab complex. (B) Representative cryo-EM micrograph and 2D averages of the FEX-CA/10E8v4 Fab complex in DDM micelles, with a 100 Å scale bar indicated. (C) Fourier shell correlation (FSC) curves as a function of resolution calculated between the half maps. (D) Local resolution map colored by local resolution estimate (B factor -137.8) using cryoSPARC. (E) Particle angular distribution plot.

**Supplementary Figure 2.**
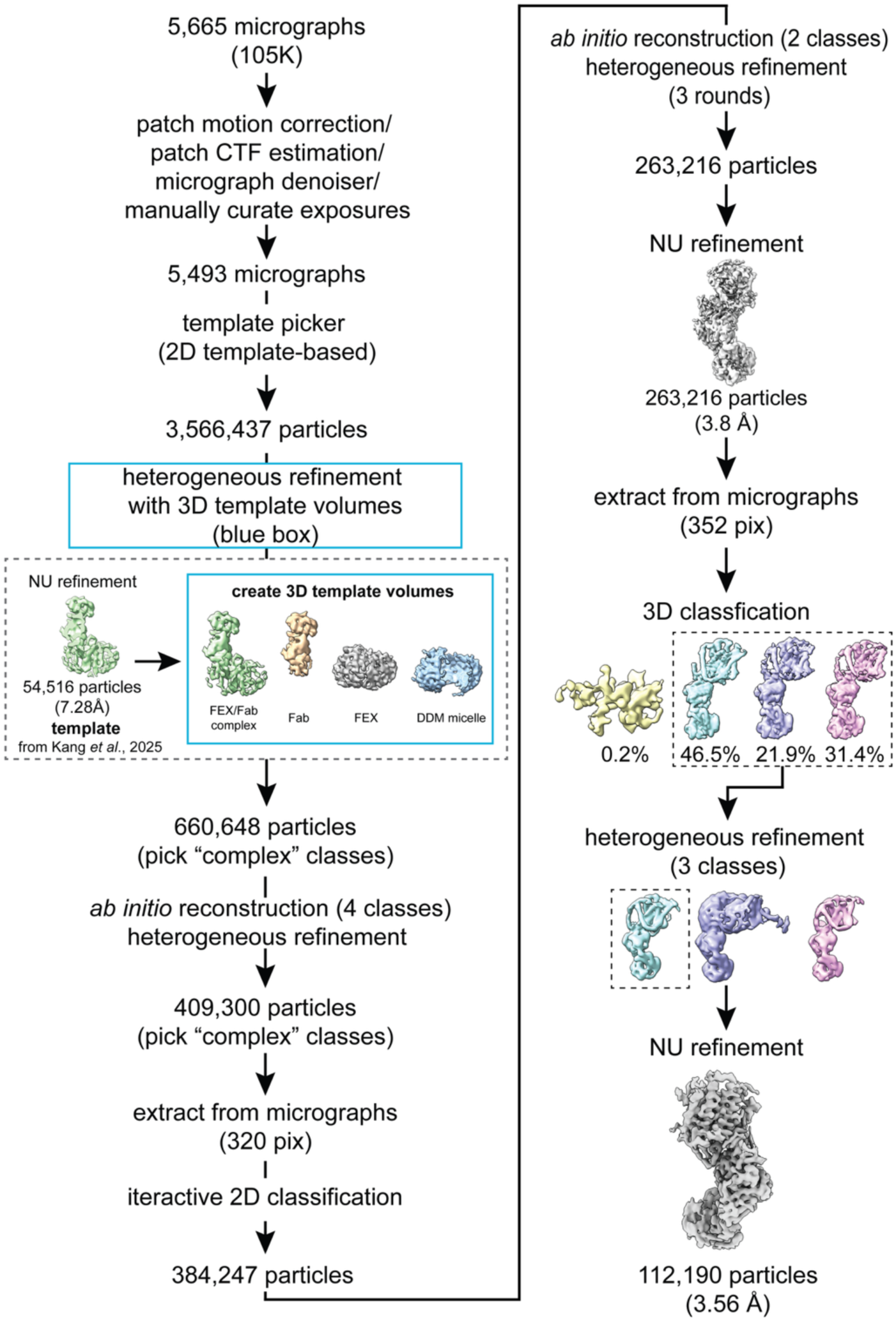
Cryo-EM workflow for data processing for FEX-CA in LiCl. All steps were processed using Cryosparc v4. 3D template volumes (blue box) were generated as described previously(9) and used for particle picking. Non-uniform refinement of the final class, containing 112,190 particles, resulted in a 3.56 Å resolution map.

**Supplementary Figure 3.**
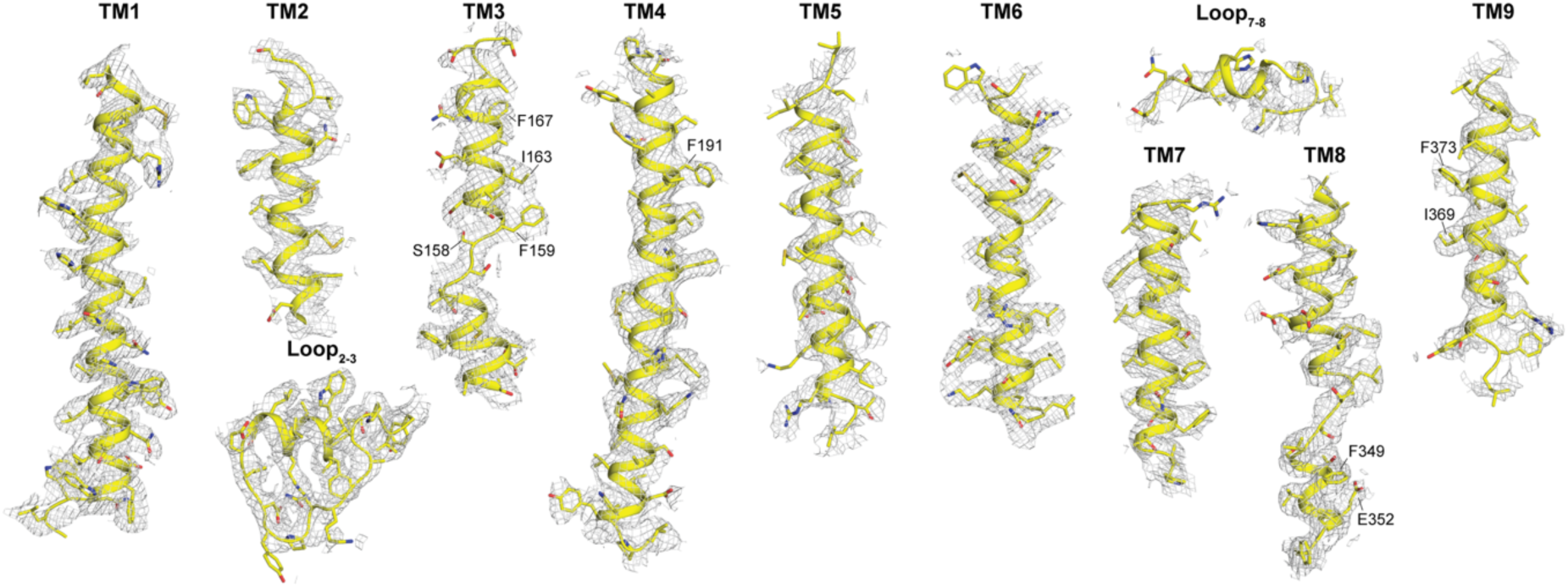
Cryo-EM maps and models of transmembrane helices for Li^+^-FEX-CA in LiCl solution. Map contoured at 6σ, with some key residues indicated.

**Supplementary Figure 4.**
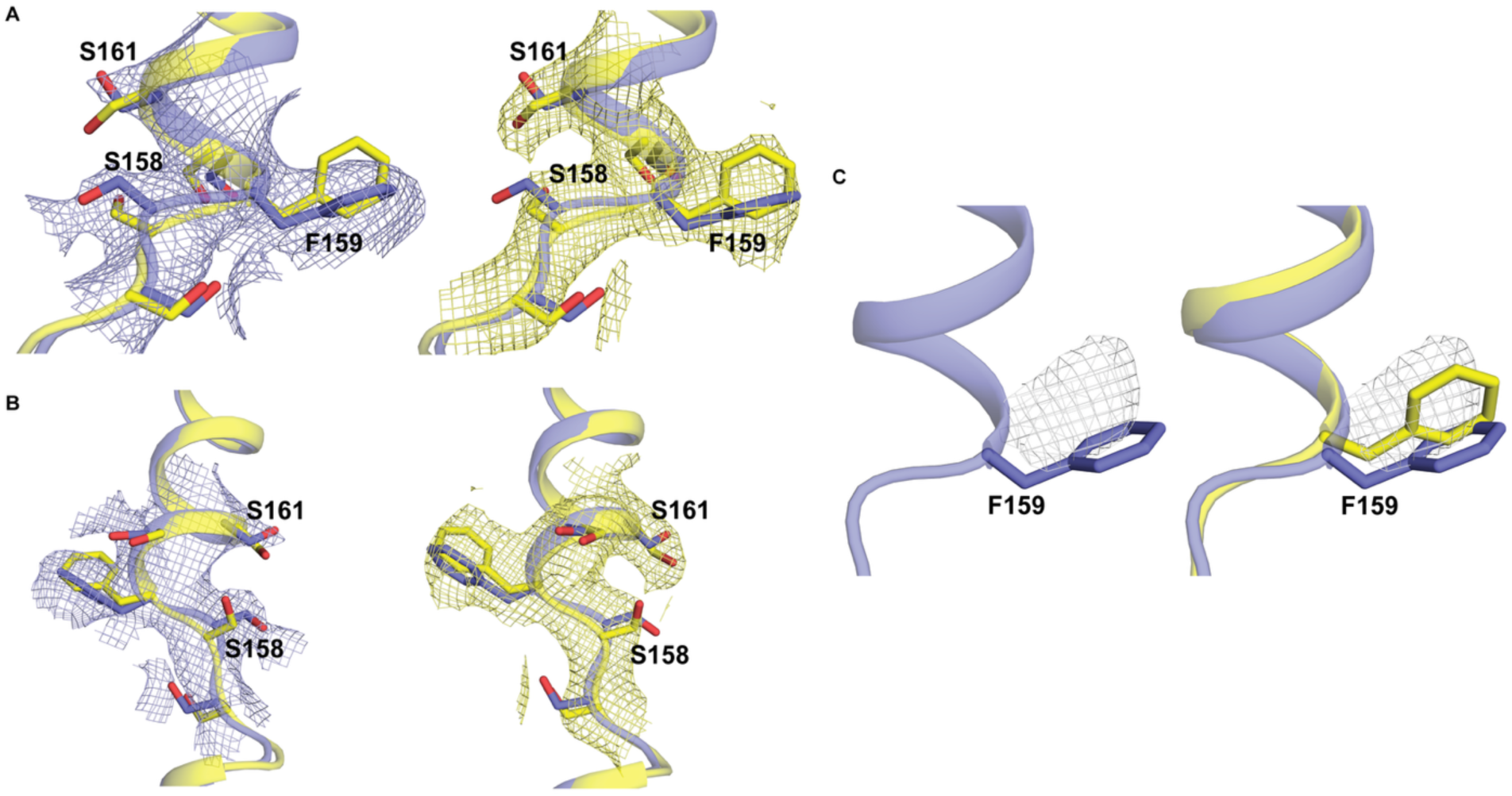
Maps and models of TM3 break in the Li^+^ and Na^+^-FEX-CA structures. (A, B) Two views of overlaid TM_3a-3b_ from the Li⁺- and Na⁺-FEX-CA models (yellow and purple, respectively). The conserved helical break (S_157_SFSS_161_) sequence is shown with stick representation. The maps for Li^+^-FEX-CA (left, yellow) and Na^+^-FEX-CA (right, purple) surrounding this sequence are contoured at 9α. (C) Difference map (gray, contoured at 13α) in the vicinity of F159 was generated by subtracting the Na^+^-FEX-CA map from the Li^+^-FEX-CA map, with both maps sharpened to the same B-factor (−137.8). Models are shown for Na^+^-FEX-CA alone (left, purple) and overlaid with Li⁺-FEX-CA (right, yellow).

**Supplementary Figure 5.**
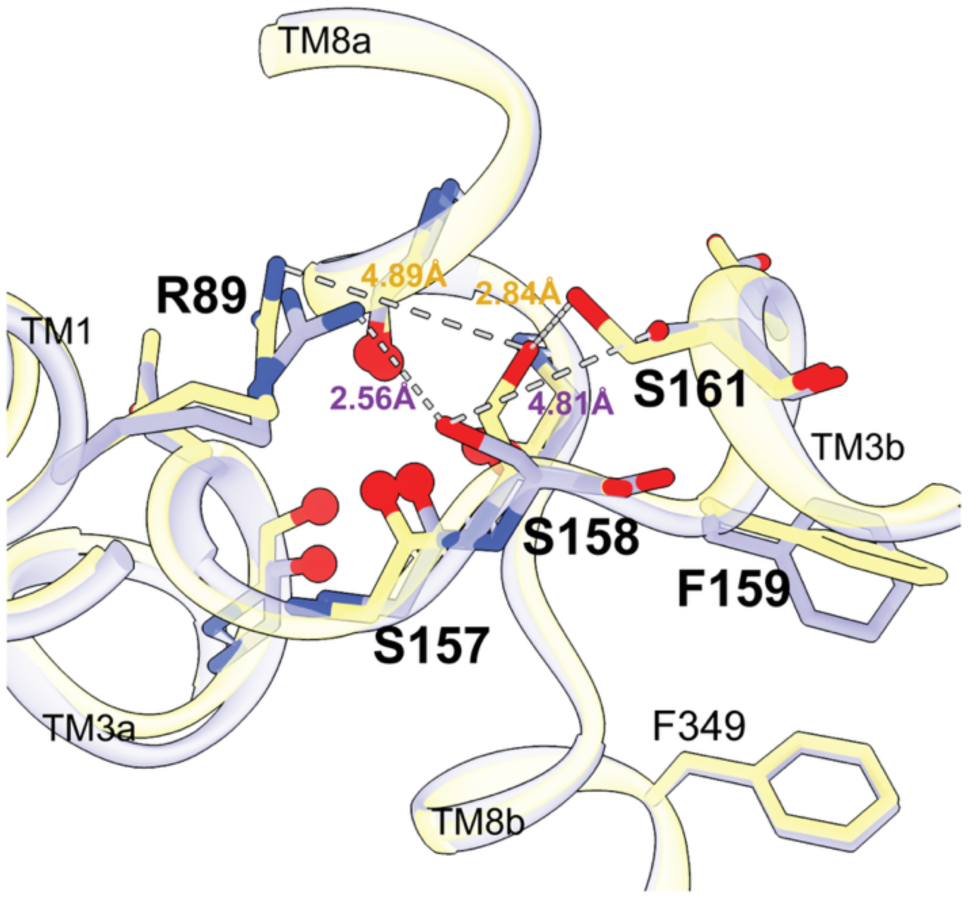
Structural comparison of Na^+^-binding site of Li^+^-FEX-CA (yellow) and Na^+^-FEX-CA (purple). The distances between S158 and R89, and between S158 and S161 are indicated by dashed lines.

**Supplementary Figure 6.**
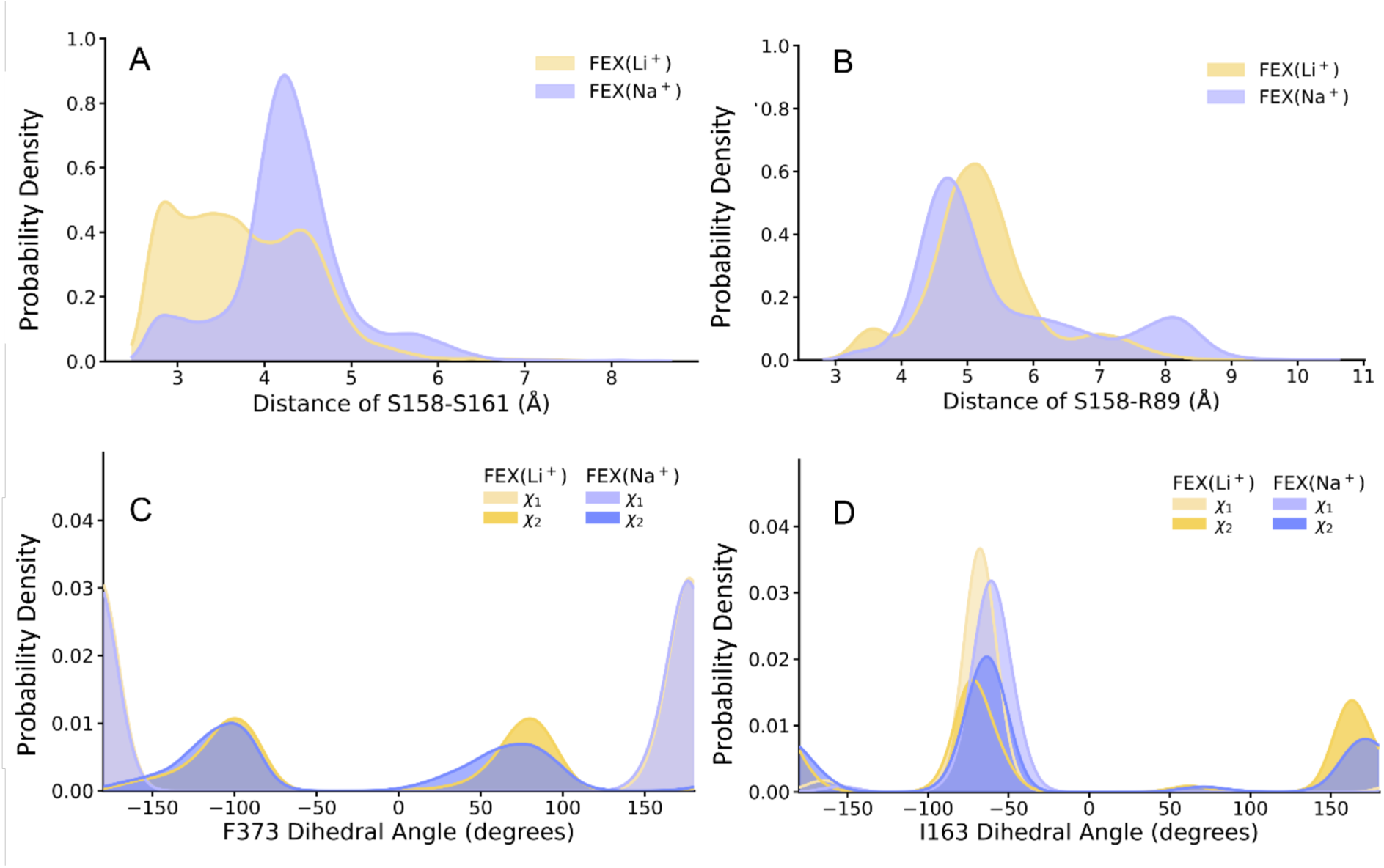
Distance and dihedral angle comparisons of selected residues in Li⁺-FEX-CA and Na⁺-FEX-CA over the course of the simulations. Probability density distributions of the distances between (A) S158 and S161 and (B) S158 and R89 in Li⁺-FEX-CA and Na⁺-FEX-CA. Probability density distributions of the χ1 and χ2 dihedral angles of (C) F373 and (D) I163 throughout the simulations.

**Supplementary Figure 7.**
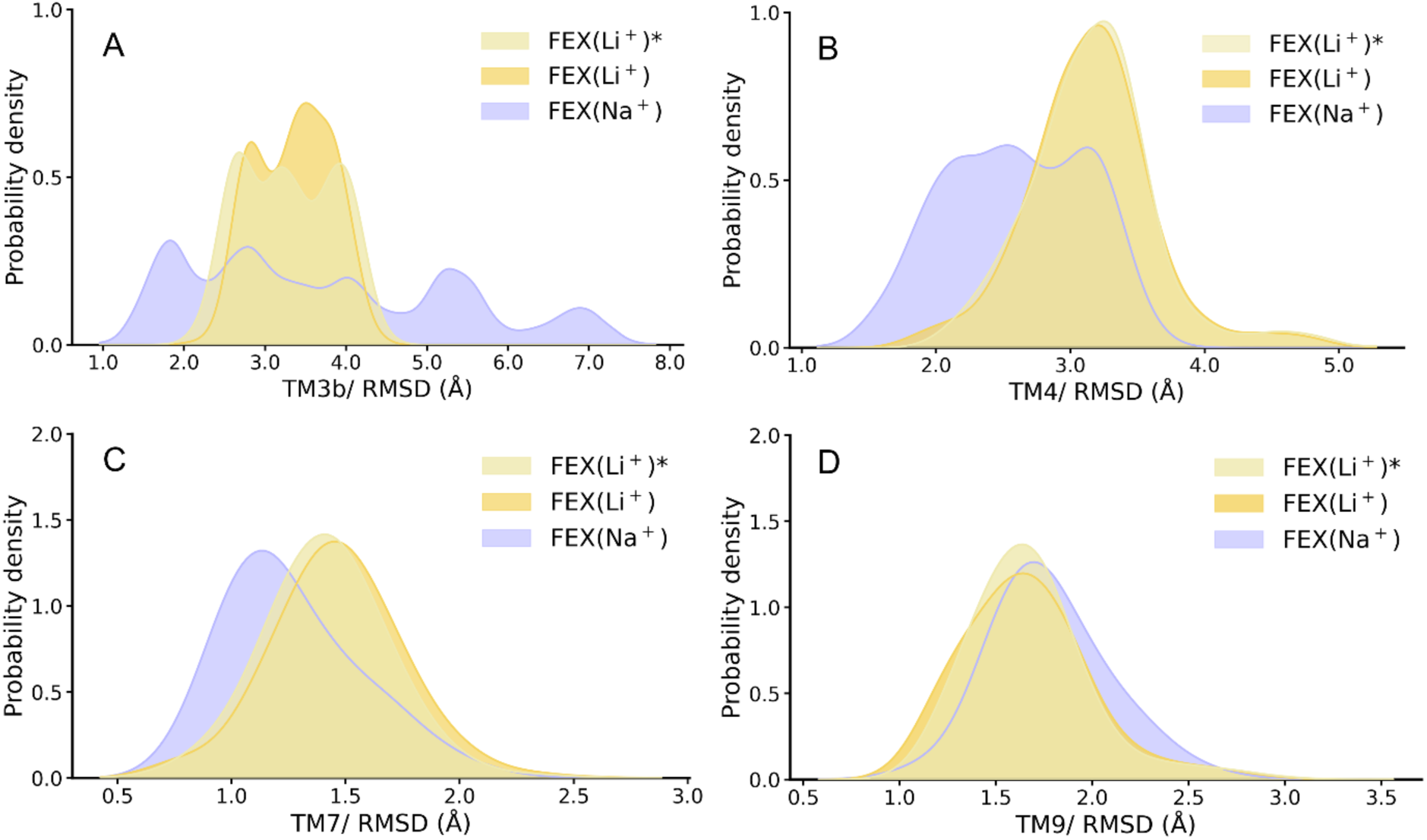
RMSD comparison of pore-lining helices in Li⁺-FEX-CA and Na⁺-FEX-CA over the simulation trajectories. Probability density distributions of RMSD for (A) TM3b, (B) TM4, (C) TM7, and (D) TM9 in Li⁺-FEX-CA and Na⁺-FEX-CA relative to the Na^+^-FEX-CA cryo-EM structures. Asterisks (*) indicate the Li⁺-FEX-CA cryo-EM structure used as the reference.

**Supplementary Figure 8.**
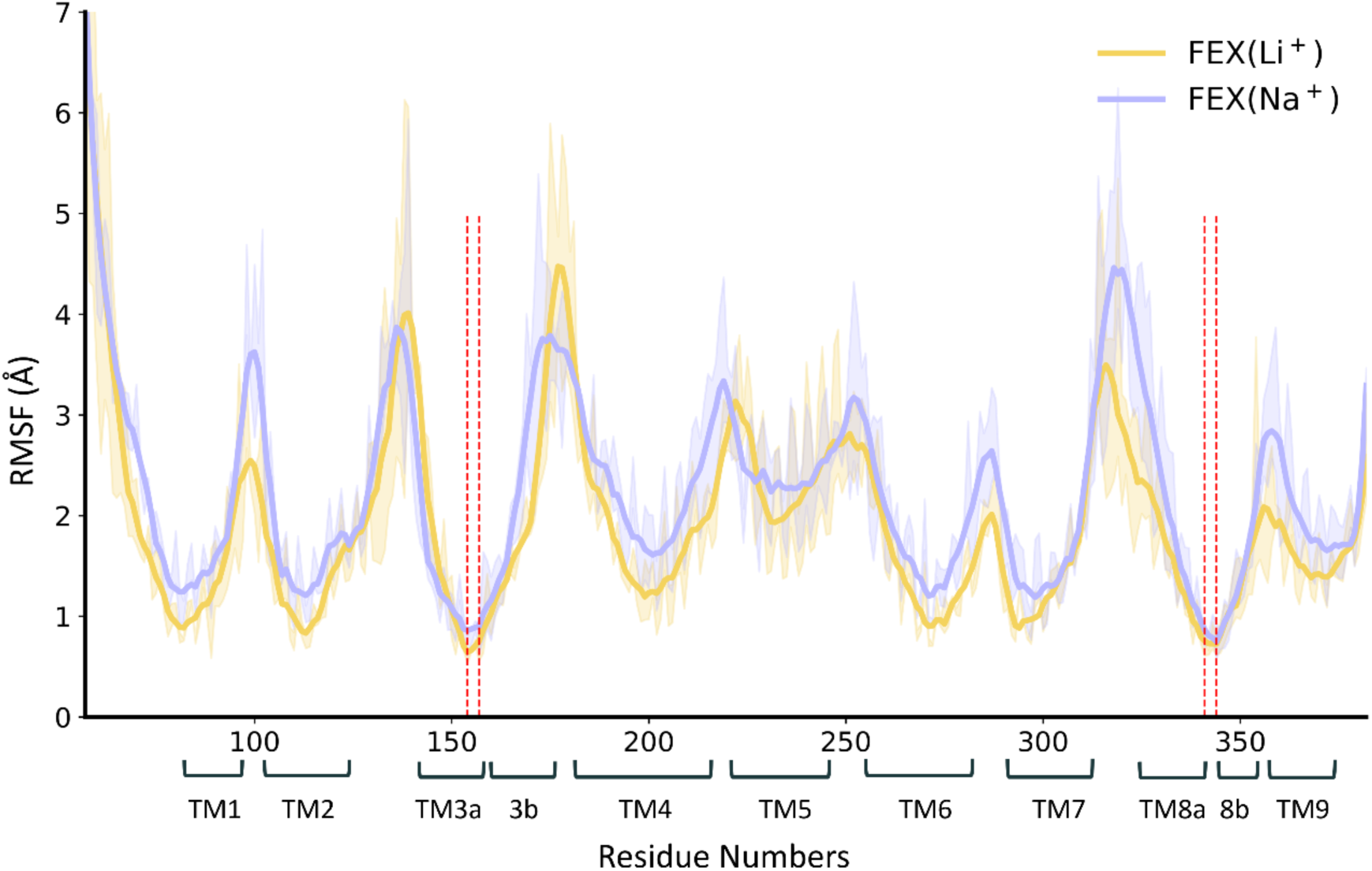
RMSF comparison of Li⁺-FEX-CA and Na⁺-FEX-CA over the simulation trajectories. Mean RMSF as a function of FEX-CA residue number, shown as the mean (solid line) ± s.d. (shaded area) across three individual replicates. The Li⁺ and Na⁺ coordinating residues G154, S157, G341, and T344 are indicated by red dashed lines.

**Supplementary Table 1.** Cryo-EM data collection, refinement, and validation statistics for FEX-CA in LiCl.

|  | FEX-CA<br>(EMD-76140)<br>(PDB 11WR) |
| --- | --- |
| **Data collection and processing** |  |
| Magnification | 105,000 x |
| Voltage (kV) | 300 |
| Bioquantum slit width(eV) | 20 |
| Electron exposure (e-/Å<sup>2</sup>) | 50 |
| Defocus range (μm) | -0.8 to -2.5 |
| Pixel size (Å) | 0.834 |
| Symmetry imposed | C1 symmetry |
| Initial particle images (no.) |  |
| Final particle images (no.) | 112,190 |
| Map resolution (Å) | 3.56 |
| Map resolution range (Å) | 3.0 to 8.1 |
| FSC threshold | 0.143 |
| **Refinement** |  |
| Initial model used | PDB 9OP1 |
| Model resolution (Å) | 3.49 |
| FSC threshold | 0.143 |
| Map sharpening B factor (Å<sup>2</sup>) | 137.8 |
| Model-to-map, CC<sub>mask</sub> | 0.81 |
| Model composition |  |
| Non-hydrogen atoms | 5,843 |
| Protein residues | 766 |
| Water | - |
| B factor (Å<sup>2</sup>) |  |
| Protein | 77.51 |
| Water | - |
| R.m.s. deviations |  |
| Bond lengths (Å) | 0.004 |
| Bond angles (°) | 0.683 |
| **Validation** |  |
| MolProbity score | 2.42 |
| Clashscore | 11.51 |
| Poor rotamers (%) | 3.3 |
| Ramachandran plot |  |
| Favored (%) | 92.5 |
| Allowed (%) | 7.37 |
| Outliers (%) | 0.13 |

## References

1. L. H. Weinstein, A. Davison, Fluorides in the environment: effects on plants and animals (CABI Publishing, Cambridge, MA, 2004).

2. R. B. Stockbridge, L. P. Wackett, The link between ancient microbial fluoride resistance mechanisms and bioengineering organofluorine degradation or synthesis. Nature Communications 15, 4593 (2024).

3. S. A. Strobel, N. R. Johnston, G. Cline, Cells Adapt to Resist Fluoride through Metabolic Deactivation and Intracellular Acidification. Chem Res Toxicol 35, 2085–2096 (2022).

4. B. C. McIlwain, M. T. Ruprecht, R. B. Stockbridge, Membrane Exporters of Fluoride Ion. Annu Rev Biochem 90, 559–579 (2021).

5. S. Li et al., Eukaryotic resistance to fluoride toxicity mediated by a widespread family of fluoride export proteins. Proceedings of the National Academy of Sciences 110, 19018–19023 (2013).

6. J. L. Baker et al., Widespread genetic switches and toxicity resistance proteins for fluoride. Science 335, 233–235 (2012).

7. S. L. Tausta, T. Berbasova, M. Peverelli, S. A. Strobel, The fluoride transporter FLUORIDE EXPORTER (FEX) is the major mechanism of tolerance to fluoride toxicity in plants. Plant Physiol 10.1093/plphys/kiab131 (2021).

8. R. B. Stockbridge, J. L. Robertson, L. Kolmakova-Partensky, C. Miller, A family of fluoride-specific ion channels with dual-topology architecture. Elife 2, e01084 (2013).

9. C. Y. Kang et al., The molecular mechanism of fluoride export by the eukaryotic fluoride channel FEX. Nat Commun 17, 589 (2025).

10. R. B. Stockbridge, A. Koide, C. Miller, S. Koide, Proof of dual-topology architecture of Fluc F-channels with monobody blockers. Nat Commun 5, 5120 (2014).

11. K. D. Smith et al., Yeast Fex1p Is a Constitutively Expressed Fluoride Channel with Functional Asymmetry of Its Two Homologous Domains. Journal of Biological Chemistry 290, 19874–19887 (2015).

12. R. B. Stockbridge et al., Crystal structures of a double-barrelled fluoride ion channel. Nature 525, 548–551 (2015).

13. B. C. McIlwain, K. Martin, E. A. Hayter, R. B. Stockbridge, An Interfacial Sodium Ion is an Essential Structural Feature of Fluc Family Fluoride Channels. J Mol Biol 10.1016/j.jmb.2020.01.007 (2020).

14. M. Ernst, E. A. Orabi, R. B. Stockbridge, J. D. Faraldo-Gomez, J. L. Robertson, Dimerization mechanism of an inverted-topology ion channel in membranes. Proc Natl Acad Sci U S A 120, e2308454120 (2023).

15. F. M. Ortega, K. Mills, H. Torabifard, Asymmetry-induced distinct mechanisms and the transporting role of sodium in bacterial fluoride channel Fluc. *bioRxiv* 10.64898/2026.02.18.706662 (2026).

16. R. B. Stockbridge, The application of Poisson distribution statistics in ion channel reconstitution to determine oligomeric architecture. Methods Enzymol 652, 321–340 (2021).

17. D. J. Wyllie, P. E. Chen, Taking the time to study competitive antagonism. Br J Pharmacol 150, 541–551 (2007).

18. P. Reith et al., The Effect of Lithium on the Budding Yeast Saccharomyces cerevisiae upon Stress Adaptation. Microorganisms 10 (2022).

19. B. C. McIlwain et al., N-terminal Transmembrane-Helix Epitope Tag for X-ray Crystallography and Electron Microscopy of Small Membrane Proteins. J Mol Biol 433, 166909 (2021).

20. T. Racek et al., MOLEonline: a web-based tool for analysing channels, tunnels, and pores (2025 update). Bioinformatics 41 (2025).

21. N. B. Last, S. Sun, M. C. Pham, C. Miller, Molecular determinants of permeation in a fluoride-specific ion channel. Elife 6 (2017).

22. T. Berbasova et al., Fluoride export (FEX) proteins from fungi, plants and animals are ’single barreled’ channels containing one functional and one vestigial ion pore. PLoS One 12, e0177096 (2017).

23. E. Chovancova et al., CAVER 3.0: a tool for the analysis of transport pathways in dynamic protein structures. PLoS Comput Biol 8, e1002708 (2012).

24. A. M. Pajor, B. A. Hirayama, D. D. Loo, Sodium and lithium interactions with the Na+/Dicarboxylate cotransporter. J Biol Chem 273, 18923–18929 (1998).

25. K. Inoue, L. Zhuang, D. M. Maddox, S. B. Smith, V. Ganapathy, Human sodium-coupled citrate transporter, the orthologue of Drosophila Indy, as a novel target for lithium action. Biochem J 374, 21–26 (2003).

26. O. Andrini, A. K. Meinild, C. Ghezzi, H. Murer, I. C. Forster, Lithium interactions with Na+-coupled inorganic phosphate cotransporters: insights into the mechanism of sequential cation binding. Am J Physiol Cell Physiol 302, C539–554 (2012).

27. E. Gopal et al., Species-specific influence of lithium on the activity of SLC13A5 (NaCT): lithium-induced activation is specific for the transporter in primates. J Pharmacol Exp Ther 353, 17–26 (2015).

28. A.-M. Sapse, P. v. R. Schleyer, Lithium chemistry : a theoretical and experimental overview (Wiley, New York, 1995), pp. x, 595 p.

29. N. Shakoor et al., Environment relevant concentrations of lithium influence soybean development via metabolic reprogramming. J Hazard Mater 441 (2023).

30. T. Dudev, K. Mazmanian, C. Lim, Competition between Li(+) and Na(+) in sodium transporters and receptors: Which Na(+)-Binding sites are “therapeutic” Li(+) targets? Chem Sci 9, 4093–4103 (2018).

31. V. Balaram, M. Santosh, M. Satyanarayanan, N. Srinivas, H. Gupta, Lithium: A review of applications, occurrence, exploration, extraction, recycling, analysis, and environmental impact. Geosci Front 15 (2024).

32. L. A. Kszos, J. J. Beauchamp, A. J. Stewart, Toxicity of lithium to three freshwater organisms and the antagonistic effect of sodium. Ecotoxicology 12, 427–437 (2003).

33. J. D. Meisel, D. H. Kim, Inhibition of Lithium-Sensitive Phosphatase BPNT-1 Causes Selective Neuronal Dysfunction in C. elegans. Curr Biol 26, 1922–1928 (2016).

34. C. A. Masuda, M. A. Xavier, K. A. Mattos, A. Galina, M. Montero-Lomeli, Phosphoglucomutase is an in vivo lithium target in yeast. J Biol Chem 276, 37794–37801 (2001).

35. L. F. Martins et al., Lithium-mediated suppression of morphogenesis and growth in Candida albicans. FEMS Yeast Res 8, 615–621 (2008).

36. K. Lai, S. D. Langley, F. W. Khwaja, E. W. Schmitt, L. J. Elsas, GALT deficiency causes UDP-hexose deficit in human galactosemic cells. Glycobiology 13, 285–294 (2003).

37. T. J. Mulkey, Alteration of growth and gravitropic response of maize roots by lithium. Gravit Space Biol Bull 18, 119–120 (2005).

38. E. A. Winzeler et al., Functional characterization of the S. cerevisiae genome by gene deletion and parallel analysis. Science 285, 901–906 (1999).

39. G. P. Lin-Cereghino et al., The effect of alpha-mating factor secretion signal mutations on recombinant protein expression in Pichia pastoris. Gene 519, 311–317 (2013).

40. C. Y. Kang, M. An, R. B. Stockbridge, Lanthanum-fluoride electrode-based methods to monitor fluoride transport in cells and reconstituted lipid vesicles. Methods Enzymol 696, 43–63 (2024).

41. Y. D. Kwon et al., Optimization of the Solubility of HIV-1-Neutralizing Antibody 10E8 through Somatic Variation and Structure-Based Design. J Virol 90, 5899–5914 (2016).

42. D. N. Mastronarde, Automated electron microscope tomography using robust prediction of specimen movements. J Struct Biol 152, 36–51 (2005).

43. A. Punjani, J. L. Rubinstein, D. J. Fleet, M. A. Brubaker, cryoSPARC: algorithms for rapid unsupervised cryo-EM structure determination. Nat Methods 14, 290–296 (2017).

44. A. Punjani, H. Zhang, D. J. Fleet, Non-uniform refinement: adaptive regularization improves single-particle cryo-EM reconstruction. Nat Methods 17, 1214–1221 (2020).

45. P. Emsley, B. Lohkamp, W. G. Scott, K. Cowtan, Features and development of Coot. Acta Crystallogr D Biol Crystallogr 66, 486–501 (2010).

46. P. V. Afonine et al., Real-space refinement in PHENIX for cryo-EM and crystallography. Acta Crystallogr D Struct Biol 74, 531–544 (2018).

47. M. G. Prisant, C. J. Williams, V. B. Chen, J. S. Richardson, D. C. Richardson, New tools in MolProbity validation: CaBLAM for CryoEM backbone, UnDowser to rethink “waters,” and NGL Viewer to recapture online 3D graphics. Protein Sci 29, 315–329 (2020).

48. E. F. Pettersen et al., UCSF ChimeraX: Structure visualization for researchers, educators, and developers. Protein Sci 30, 70–82 (2021).

49. R. Anandakrishnan, B. Aguilar, A. V. Onufriev, H++ 3.0: automating pK prediction and the preparation of biomolecular structures for atomistic molecular modeling and simulations. Nucleic Acids Res 40, W537–541 (2012).

50. J. C. Gordon et al., H++: a server for estimating pKas and adding missing hydrogens to macromolecules. Nucleic Acids Res 33, W368–371 (2005).

51. W. L. Jorgensen, J. Chandrasekhar, J. D. Madura, R. W. Impey, M. L. Klein, Comparison of Simple Potential Functions for Simulating Liquid Water. J Chem Phys 79, 926–935 (1983).

52. S. Schott-Verdugo, H. Gohlke, PACKMOL-Memgen: A Simple-To-Use, Generalized Workflow for Membrane-Protein-Lipid-Bilayer System Building. J Chem Inf Model 59, 2522–2528 (2019).

53. D. A. Case et al., AMBER 2020 (University of California, San Francisco, 2020).

54. J. A. Maier et al., ff14SB: Improving the Accuracy of Protein Side Chain and Backbone Parameters from ff99SB. J Chem Theory Comput 11, 3696–3713 (2015).

55. C. J. Dickson, R. C. Walker, I. R. Gould, Lipid21: Complex Lipid Membrane Simulations with AMBER. J Chem Theory Comput 18, 1726–1736 (2022).

56. A. W. Gotz et al., Routine Microsecond Molecular Dynamics Simulations with AMBER on GPUs. 1. Generalized Born. J Chem Theory Comput 8, 1542–1555 (2012).

57. R. J. Loncharich, B. R. Brooks, R. W. Pastor, Langevin dynamics of peptides: the frictional dependence of isomerization rates of N-acetylalanyl-N’-methylamide. Biopolymers 32, 523–535 (1992).

58. H. J. C. Berendsen, J. P. M. Postma, W. F. Vangunsteren, A. Dinola, J. R. Haak, Molecular-Dynamics with Coupling to an External Bath. J Chem Phys 81, 3684–3690 (1984).

59. J. P. Ryckaert, G. Ciccotti, H. J. C. Berendsen, Numerical integration of the cartesian equations of motion of a system with constraints: molecular dynamics of n-alkanes. Journal of computational physics 23, 327–341 (1977).

60. D. R. Roe, T. E. Cheatham, 3rd, PTRAJ and CPPTRAJ: Software for Processing and Analysis of Molecular Dynamics Trajectory Data. J Chem Theory Comput 9, 3084–3095 (2013).

61. W. Humphrey, A. Dalke, K. Schulten, VMD: visual molecular dynamics. J Mol Graph 14, 33–38, 27-38 (1996).

62. H. Yu et al., Simulating Monovalent and Divalent Ions in Aqueous Solution Using a Drude Polarizable Force Field. J Chem Theory Comput 6, 774–786 (2010).

63. W. S. Timothy Baker (2026) ResiRuler. (GitHub).

